# Sub-3 Å resolution structure of apoferritin using a multi-purpose TEM with a side-entry cryo-holder

**DOI:** 10.1101/2020.03.24.006619

**Authors:** Yoko Kayama, Raymond N. Burton-Smith, Chihong Song, Naoya Terahara, Takayuki Kato, Kazuyoshi Murata

## Abstract

The structural analysis of protein complexes by cryo-electron microscopy (cryo-EM) single particle analysis (SPA) has had great impact as a biophysical method in recent years. Many results of cryo-EM SPA are based on state-of-the-art cryo-electron microscopes customized for SPA. These are currently only available in limited locations around the world, where securing machine time is highly competitive. One potential solution for this time-competitive situation is to reuse existing multi-purpose equipment. Here, we used a multi-purpose TEM with a side entry cryo-holder at our facility to evaluate the potential of high-resolution SPA. We report a 3 Å resolution map of apoferritin with local resolution extending to 2.6 Å. The map clearly showed two positions of an aromatic side chain. We also verified the optimal imaging conditions depending on different electron microscope and camera combinations. This study demonstrates the possibilities of more widely available and established electron microscopes, and their applications for cryo-EM SPA.

## Introduction

Cryo-electron microscopy (cryo-EM) single particle analysis (SPA) is a technique for reconstructing the three-dimensional structure of a biomacromolecule using projected images acquired with an electron microscope (Bhella, 2019) and was the subject of the Nobel Prize for Chemistry in 2017 (Cressey and Callaway, 2017). The technique has achieved tremendous progress by integrating various technologies (Murata and Wolf, 2018).

Developments contributing to advances in SPA have been mainly improvements of electron microscope performance (Knapek et al., 1982; Morishita et al., 2013), developments of electron beam direct detectors (Bammes et al., 2012; McMullan et al., 2016), methods for three-dimensional structure reconstructions (Grant et al., 2018; Kimanius et al., 2016; Punjani et al., 2017; Zivanov et al., 2018), and automated acquisition via Leginon (Carragher et al., 2000), SerialEM (Mastronarde, 2005) or manufacturer software. In recent years, close to atomic resolution has been achieved (Bartesaghi et al., 2015; Danev et al., 2019; Kato et al., 2019) which permits construction of atomic models without foreknowledge of the protein sequence.

All the above-mentioned techniques are indispensable for improving achieved resolution. However, focusing on the performance of the electron microscope, including electron source, the sample stage, and the detector is arguably the primary limiting factor. For example, autoloader stages such as those used in Titan Krios (Thermo Fisher Scientific) and CRYOARM (JEOL) microscopes demonstrate that multiple sample grids can be stored stably for a long period of time, the sample grid can be automatically transported, and data can be automatically collected without manual intervention. Such a sample stage is difficult to introduce later into a multi-purpose electron microscope and is currently only available pre-installed. Such electron microscopes are very expensive and are currently only available at limited locations. As a result, competition for machine time is high. One solution is to reuse established equipment.

Optimisations of the microscope for SPA are often incompatible or non-ideal for other techniques for which the microscope could be used, such as electron tomography (Baumeister, 2002), EDS (Allen et al., 2012), EELS (Egerton, 2009), STEM (Crewe et al., 1970) and microED (Nannenga and Gonen, 2019). Nevertheless, the vast cost of maintaining multiple pieces of optimised equipment precludes general availability. Therefore, it is desirable to be aware of how realistic mid-to high-resolution SPA is on multi-purpose TEMs.

While datasets exceeding 1,000 micrographs are now regularly collected (Iudin et al., 2016), this is unrealistic for manual data collection. Therefore, in this study, limited datasets (<200 micrograph movies each) were manually collected with a test specimen of apoferritin on two microscope-and-detector combinations which serve as multi-purpose (S)TEMs, both using Gatan 626-type cryo-specimen holders. Apoferritin (Richter, 1959; Toussaint et al., 2007) is a relatively recent addition to the “benchmark” samples for cryo-EM, since its compact size, spherical shape and octahedral symmetry presented difficulties in reconstructing at lower resolutions (Russo and Passmore, 2014). Using a beta release of RELION 3.1 and <300 micrograph movies, a 3 Å (global) resolution map of apoferritin was achieved. We further examined optimized data collection conditions for each general purpose cryo-EM setting, although as early datasets here had been processed with RELION 3.0, we chose to continue processing these datasets with RELION 3.0, rather than introduce a further variable. The limited number of acquired micrographs is intended to offset the multiplicative effect of high symmetry on particle count and provide some indication of utility on lower symmetry datasets.

While not all multi-purpose TEMs are equipped with automation software, the decrease in workload for the microscope operator presented by automation, combined with the ability to collect data more quickly makes software-controlled data acquisition highly desirable. To demonstrate the utility of automation on data collection with a multi-purpose TEM which still requires manual cryogen maintenance, we acquired a dataset of β-galactosidase using SerialEM, post-installed automated software (Mastronarde, 2005) and processed independently. As a result, 3.6 Å resolution map was able to be acquired with one semi-automated session of data collection that took six hours with two replenishments of liquid nitrogen.

In this work we provide a possibility to reuse existing equipment for high-resolution cryo-EM and guidelines for the minimum setup for growth of the research field within general-access facilities.

## Results

### 3 Å resolution map reconstruction using a multi-purpose TEM with a side-entry cryo-holder

With a limited dataset of 279 micrograph movies and using a beta version of RELION 3.1, we achieved 3 Å (global) resolution of apoferritin (Fig. 1) when estimated at the gold-standard (GS) (fully independent half-maps) FSC (0.143) (Fig. S1A) using a combination of JEM-2100F electron microscope and K2 Summit direct electron detector (DED). Local resolution estimated by the *blocres* module of Bsoft (Heymann, 2001; Heymann and Belnap, 2007) shows significant areas of the map between 2.6-2.8 Å (Fig. 1A). All helices were clearly defined (Fig. 1A, B) with some residues exhibiting two conformational states (Fig. 1B, marked with black arrows) although with one conformation dominant as the second was lost at higher map σ (Fig. S1B). In higher resolution regions side chains are clear (Fig. 1C) and densities could be assigned to metal atoms coordinated by side chains (Fig. 1B, 1C, marked with red arrow). It may be possible to assign water to some densities, but we erred on the side of caution with respect to interpreting potential water-related density.

**Figure 1.**
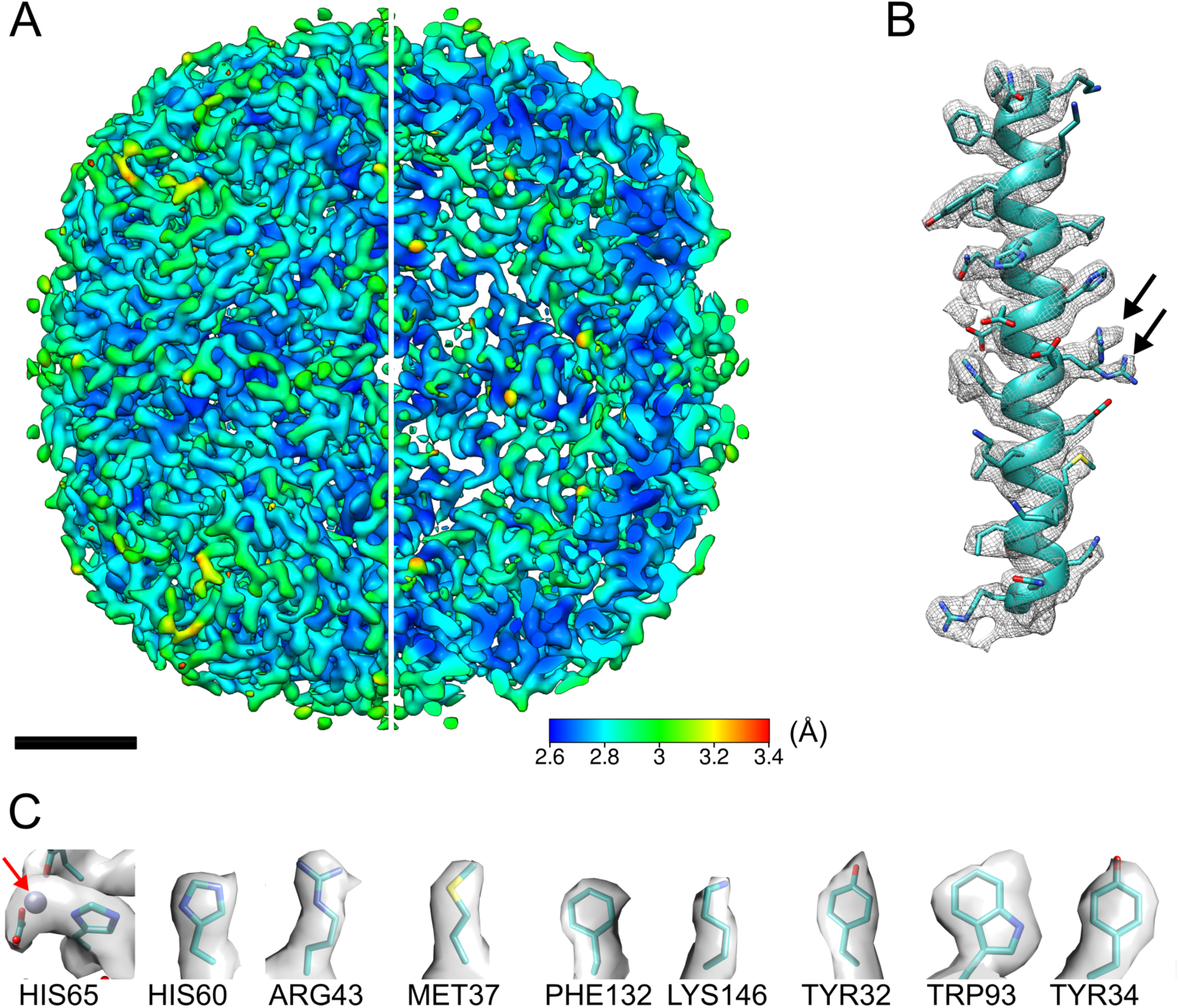
3 Å (global estimated resolution) cryo-EM map of mouse heavy-chain apoferritin. A) Map coloured by local resolution; one quarter sliced away to allow visualisation of internal density, scale bar 2 nm. B) Representative helix (Leu48-Arg77) from one subunit, with PDB:2CIH fitted. Map is contoured at 3σ. Black arrows show two rotamers of Arg63. C) Nine representative sidechains. Red arrow shows a metal density.

### Imaging comparison of two DEDs

Photographs of one EM setup (Setting A; JEM-2100F, K2 Summit DED and Gatan 626 side entry cryo-specimen holder) are shown in Fig. S2. The second (Setting B; JEM-2200FS, DE-20 DED and Gatan 626 cryo-holder) has been shown previously (Murata and Wolf, 2018). Micrograph movies independently collected from the two EM setups were corrected for image drift and electron beam damage using the MotionCor2 algorithm (Zheng et al., 2017) as implemented in RELION 3 (Zivanov et al., 2018). Table 1 details the essential information for each equipment setting.

**Table 1.**
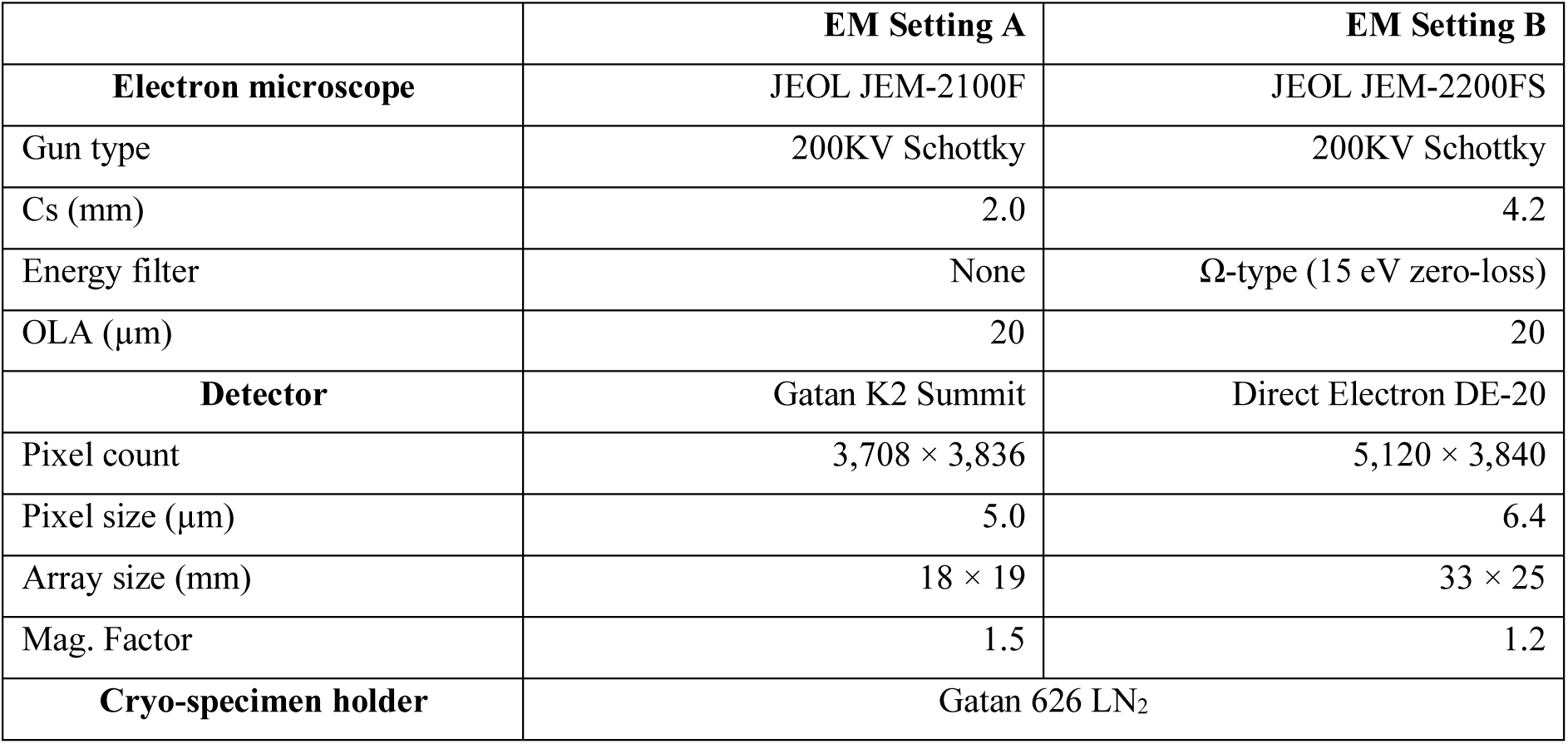
Details of TEM Settings A and B

For comparison, representative cryo-EM images from each setup are shown in Fig. 2. In this figure, images were captured at 50,000× magnification and displayed at the same particle scale. In both cases, the projected image of apoferritin particles was easily recognized with similar contrast. An area of the same absolute size as the image obtained with Setting A (K2 Summit detector) acquisition is shown on the Setting B (DE-20 detector) micrograph by a white dashed box (Fig. 2B), highlighting the difference in field of view. The total number of apoferritin particles in the example images were counted, totalling 383 in the Setting A image and 1,334 in the Setting B image (Fig. 2).

**Figure 2.**
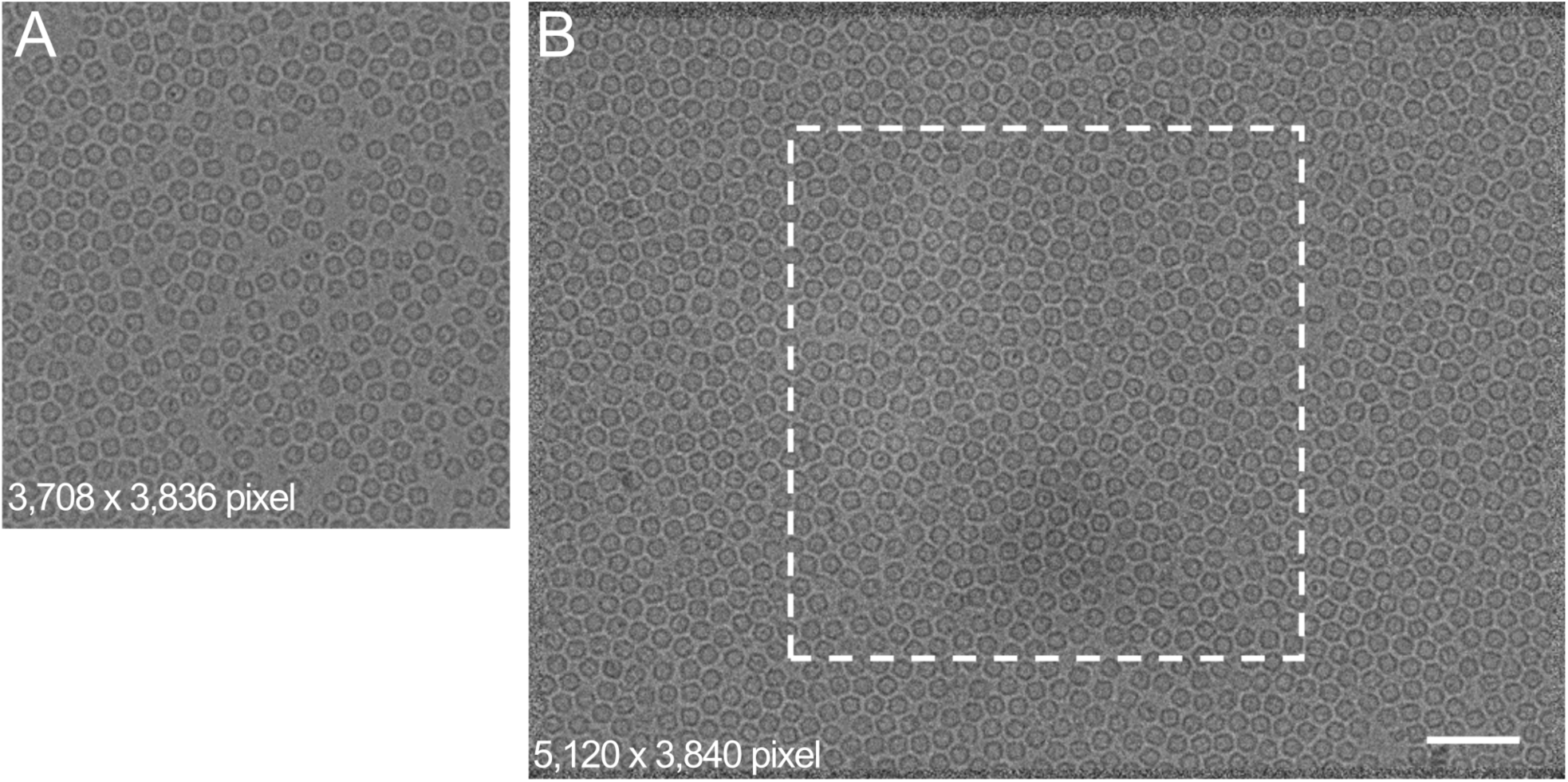
Comparison of micrographs from each microscope and detector combination. Representative micrographs from K2 Summit at 50,000× (A) and from DE-20 at 50,000× (B), each 2.4 µm under-focus. Micrographs scaled to equivalent size, scale bar 50 nm. White dashed box overlaid on image (B) highlights field of view difference between detectors. Micrograph dimensions are included in each figure. As a test sample, purified apoferritin was used. The number of particles included each micrograph were (A) 383, (B) 1,334, respectively. Absolute pixel dimensions are 5.0 µm^2^ for K2 Summit DED and 6.4 µm^2^ for DE-20 DED.

We calculated the MTF and DQE curves (Fig. S3) in each case at 200kV using a beam stopper and the FindDQE program (Ruskin et al., 2013), where the greatest difference between the curves was shown in the low frequencies. The calculated DQE value at lower special frequencies was >80% for the K2 Summit, but was reduced to ∼35% when using the DE-20. At frequencies above ¾ Nyquist, the two detectors have very similar response curves (Fig. S3). They show similar characteristics to the same DEDs on different microscopes (Faruqi and McMullan, 2018; Kuijper et al., 2015; Ruskin et al., 2013) as expected.

When comparing Setting A and Setting B using Pt-Ir film (Hamaguchi et al., 2019) (Fig. S4) the clarity of Thon rings in Setting A is immediately apparent in the power spectrum (Fig. S4A) and in the rotational average profile (Fig. S4B) to beyond the diffraction ring of 2.27 Å. Both Settings were observed at 100,000× magnification, at which point Setting B is difficult to keep stable. This is manifest in the weaker oscillations in the CTF and diffraction ring at ∼2.3 Å (Fig. S4C). Minor astigmatism of approximately 60 nm causes the blurring of the diffraction ring in Setting B upon rotational averaging, which while visible in the power spectrum is not as clear in the radial profile (Fig. S4D).

### Comparison of 3D reconstruction maps in different imaging conditions

Eight cryo-EM maps of apoferritin with resolutions between 3.3 Å and 5.1 Å were obtained by standard operation (Fig. 3) of RELION 3.0 using different data sets as detailed in Table 2. In all final post-processing steps, a very soft mask (15 Å low-pass filter, 5 pixel expansion and 10 pixel soft edge) was used, which slightly reduces the resolution estimated by “gold-standard” FSC (GS-FSC) (Chen et al., 2013) than when a less soft mask was used. We would prefer to underestimate the GS-FSC resolution than overestimate. Local resolution is unaffected by this softer mask.

**Table 2.**
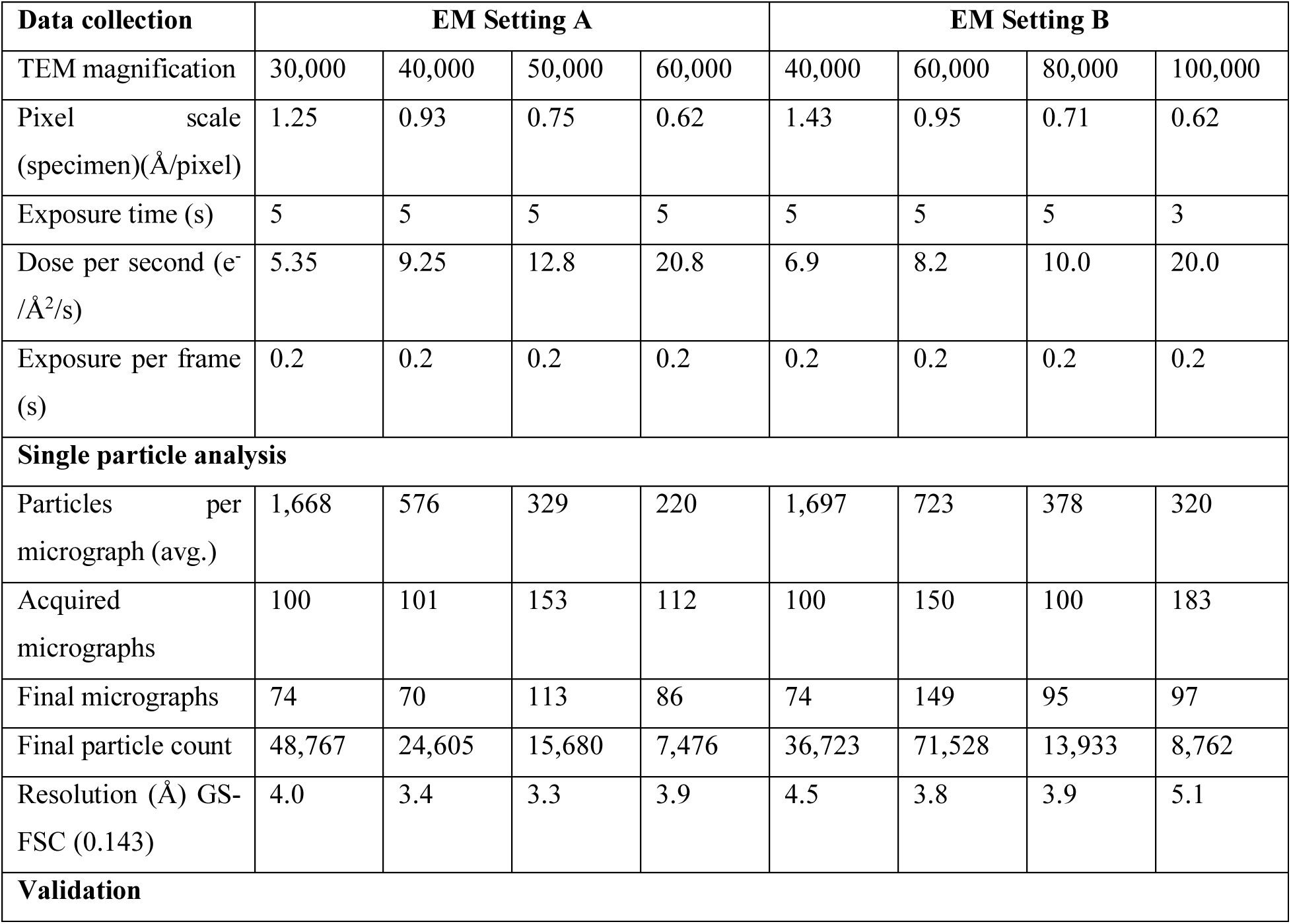
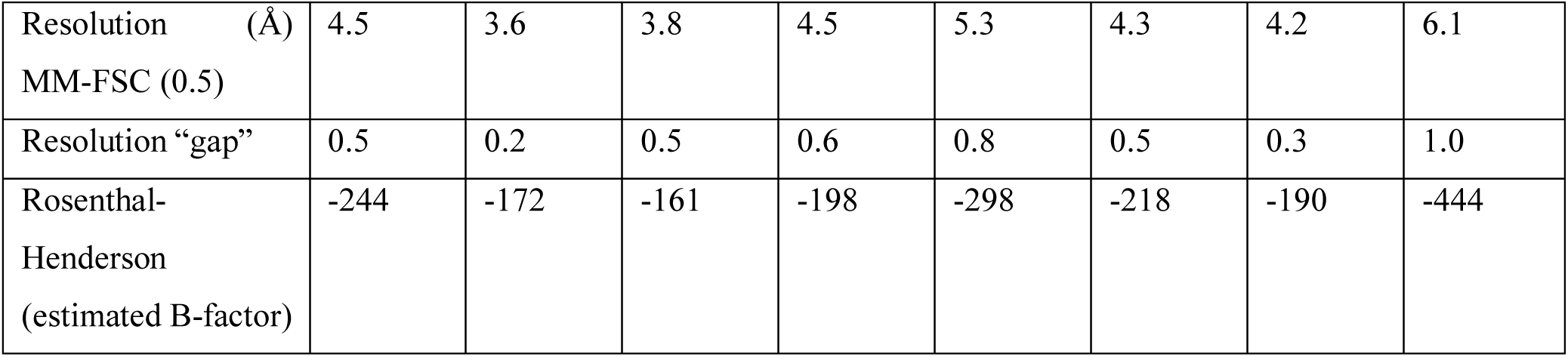
Details of each of eight acquisition conditions

**Figure 3.**
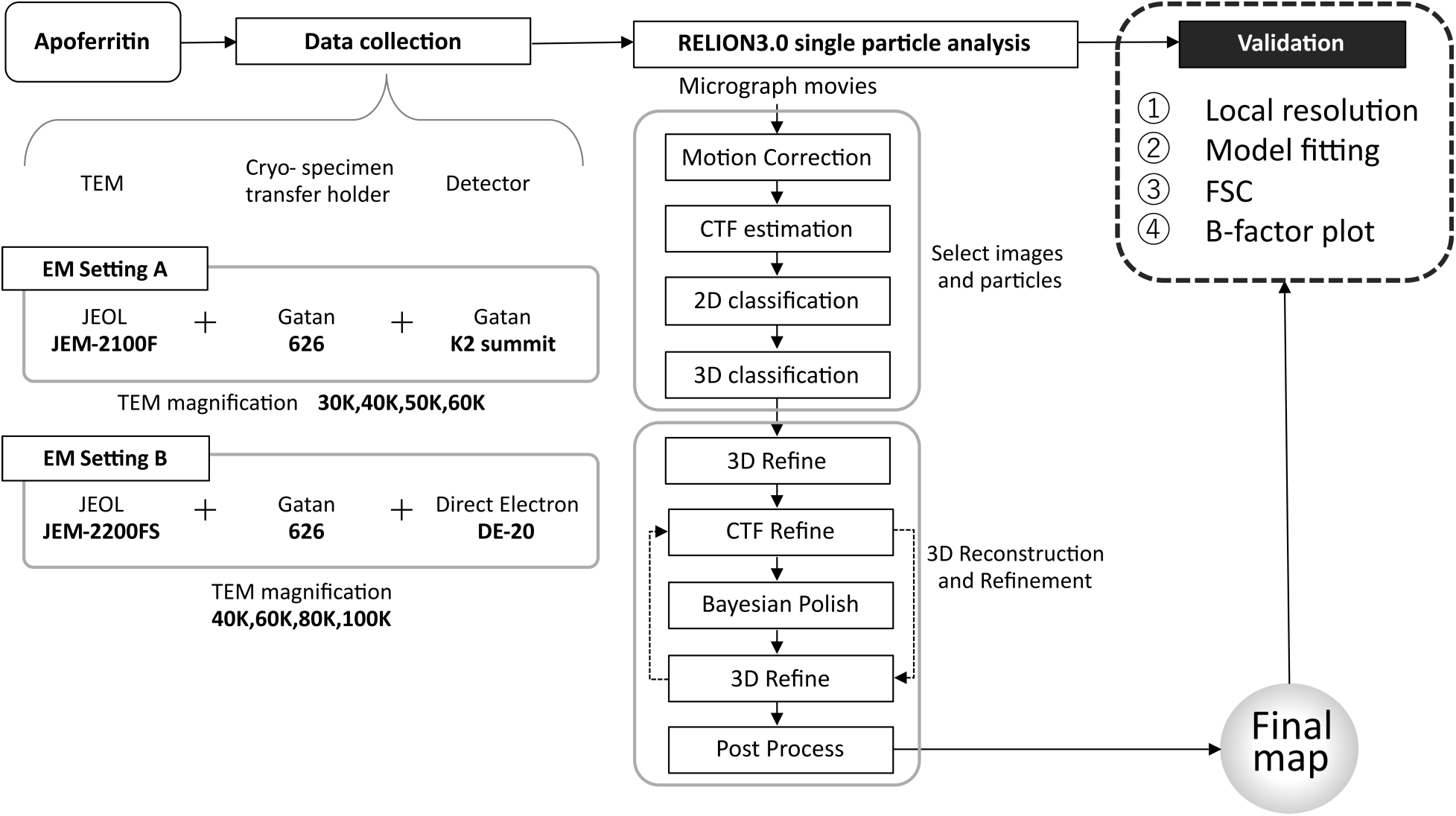
Processing workflow. General schematic of data acquisition and processing is described. Two electron microscope setups were used: JEM-2100F + Gatan 626 + Gatan K2 Summit detector (Setting A) and JEM-2200FS + Gatan 626 + Direct Electron DE-20 detector (Setting B). Each data set was processed using RELION 3.0 and evaluated with procedures shown in the figure. Further specifics of the equipment can be found in Table 1.

Using limited datasets described in Table 2, the highest resolution map was obtained from a data set acquired with Setting A at a magnification of 50,000×. The map coloured by local resolution (Heymann, 2001; Heymann and Belnap, 2007) is shown in Fig. S5. The best local resolution of ∼2.8 Å was shown in the α-helices located inside the core. On the other hand, the disordered N-terminal of each subunit showed the lowest resolution of ∼3.6 Å.

A crystal structure model (PDBID: 2CIH) (Toussaint et al., 2007) was fitted to the map and a single helix was extracted from each reconstruction to visualize map quality (Fig. 4). The highest resolution map (Fig. 4C) showed good agreement with the model to confirm the secondary structure and the side chains. Similarly, Fig. 4B also shows reasonable clarity for residue side chains. The slightly lower resolution maps (Fig. 4D, F, G, H) show moderately resolved side chains.

The best resolution map obtained using Setting B reported 3.8 Å at 60,000× magnification, but there were some areas where the electron densities of the side chains cannot be clearly recognized; lysine and arginine residues provide examples (Fig. 4, marked with black arrows).

**Figure 4.**
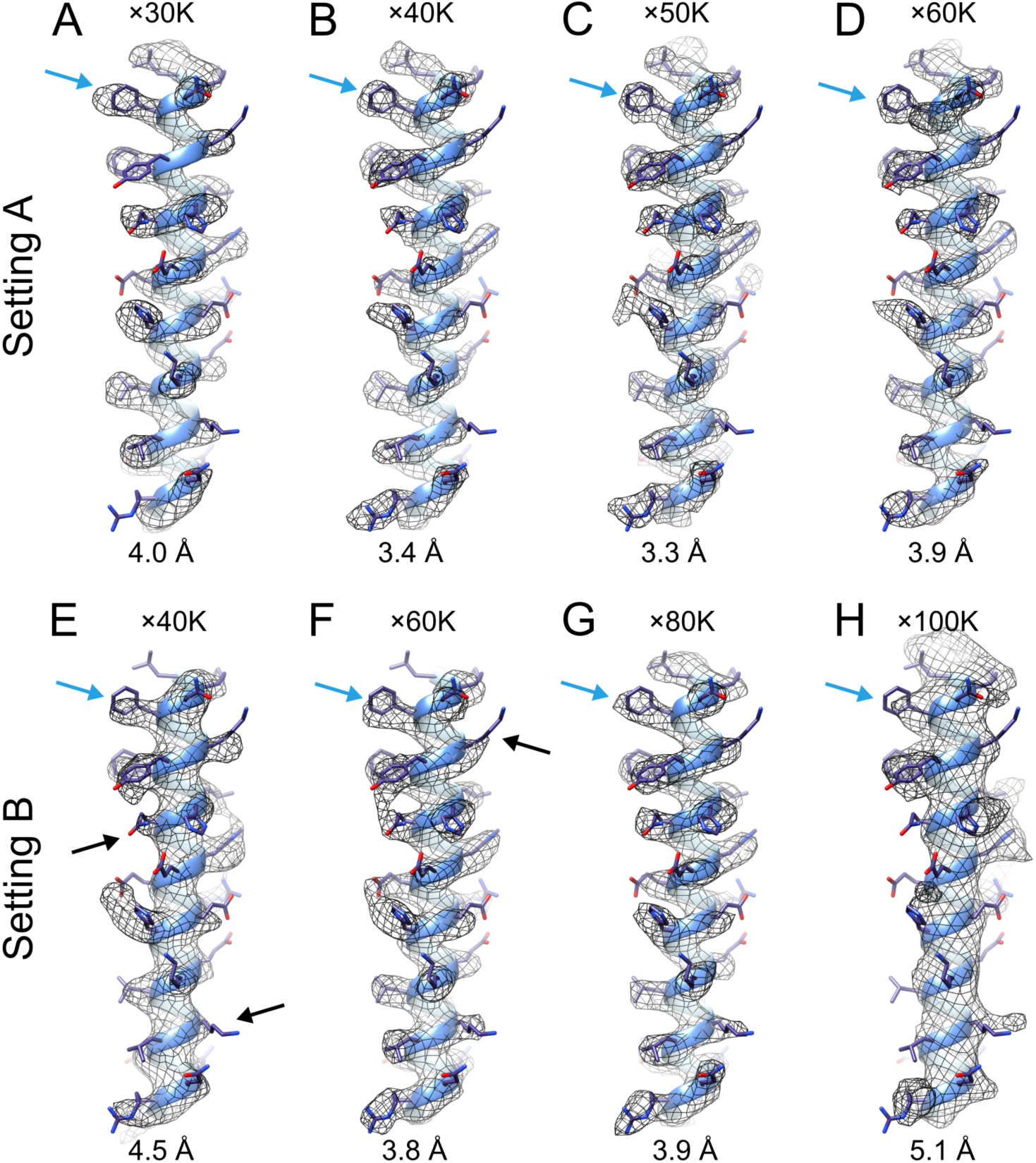
Density comparison between eight reconstruction segments. A representative α-helix (Leu48-Arg77) was extracted from each cryo-EM map and the corresponding atomic model (PDB ID: 2CIH) was fitted to each map. The maps in the top panel are reconstruct from datasets acquired by Setting A, at magnifications of ×30K (A), ×40K (B), ×50K (C), and ×60K (D). The map in a bottom panel are reconstructed from data sets acquired by Setting B, at magnifications of ×40K (E), ×60K (F), ×80K (G), and ×100K (H). The obtained resolutions are labelled in each map. Leu69 is indicated by blue arrows. Black arrows highlight exemplar sidechains which may present difficulties in identification.

While it is still possible to determine larger side chains such as tryptophan, tyrosine and phenylalanine in maps of approximately 4 Å (Fig. 4A, marked with blue arrow), it is difficult to confirm the side chains in maps worse than 4.5 Å resolution (Fig. 4E). In maps with resolutions below 5 Å (Fig. 4H), it is not possible to reliably identify side chains and unstructured loops. At 6-8 Å only α-helices were clarified (Rosenthal and Rubinstein, 2015) as demonstrated by the initial model used (Fig. S6).

### Comparison of data quality in different image conditions

In order to verify the reconstructed maps, the GS-FSC and the map-to-model FSC (MM-FSC) were compared (Fig. 5). The gap between the two FSC values were listed in Table 2. The difference values (Resolution “gap” in Table 2) were averaged in each case, showing (average ± standard error) 0.45±0.075 Å in Setting A and 0.65±0.15 Å in Setting B. These smaller gaps between GS-FSC and MM-FSC of the K2 Summit correlate with the better-quality densities as shown in the maps (Fig. 4, Table 2). This may also come from the result of the superior DQE curve (Fig. S3) and Thon rings of the K2 Summit DED (Fig. S4).

**Figure 5.**
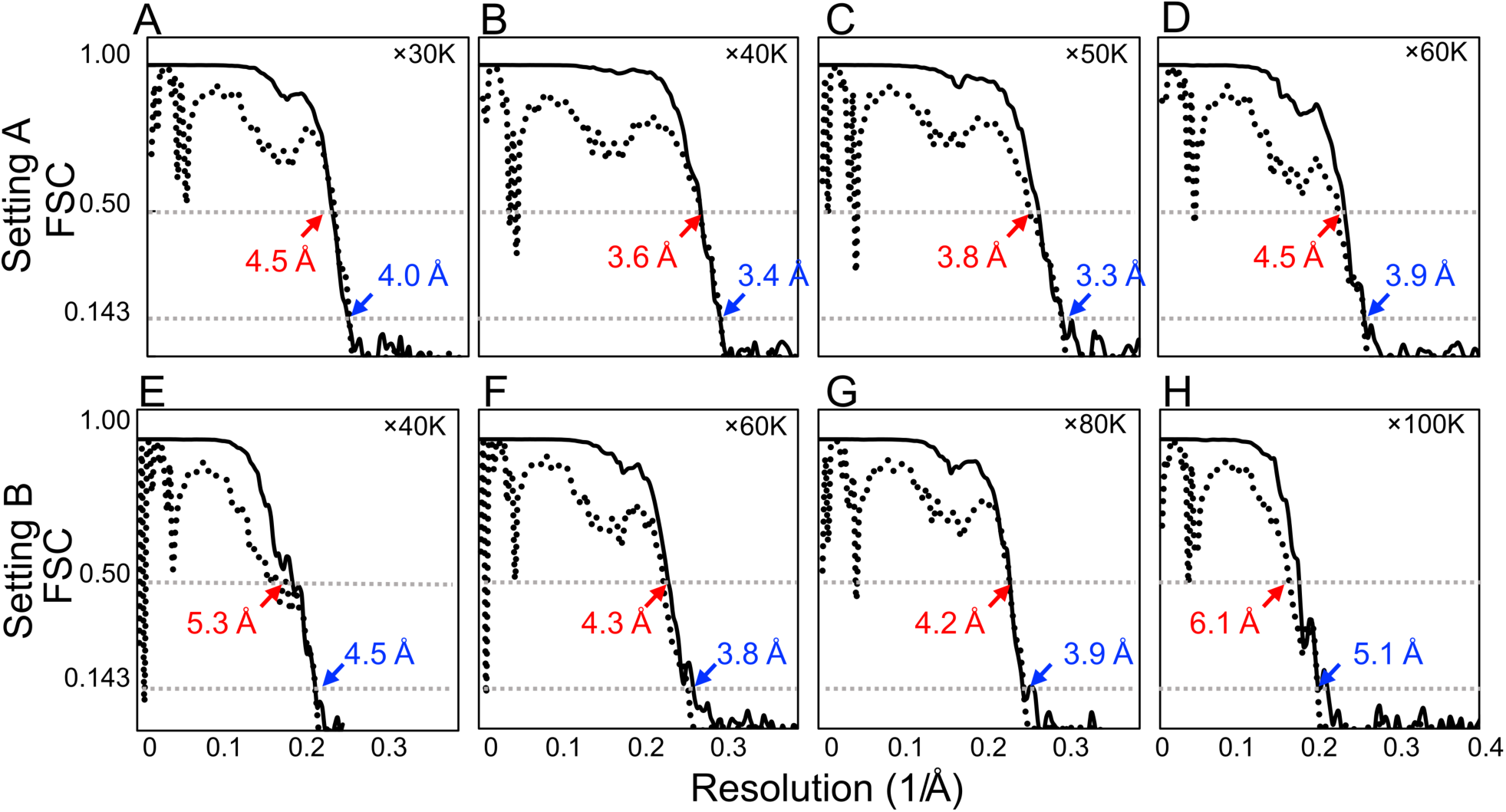
GS-FSC and MM-FSC plots of all reconstructions. GS-FSC (solid lines) was calculated from independent half maps, and the resolutions was estimated at 0.143 cut-off (blue labels). MM-FSC (dashed lines) were calculated between 3D reconstructions and maps calculated from the PDB model (2CIH), and the resolution was estimated at 0.5 cut-off (red labels). The experimental conditions are ×30K (A), ×40K (B), ×50K (C), and ×60K (D) magnifications using Setting A, and ×40K (E), ×60K (F), ×80K (G), and ×100K (H) magnifications using Setting B.

The reported resolutions of Setting A between 30,000× and 60,000× magnifications showed a similar parabola shaped curve with Setting B between 40,000× and 100,000× magnifications (Fig. 6). The sampling scales at the specimen were nearly identical within these magnification ranges (Fig. 6, Table 2). The result suggests that there is an optimal magnification for high-resolution analysis in each EM setting. The highest resolution was obtained at 50,000× magnification using Setting A, which correspond to the sample scale of 0.75 Å/pixel (Fig. 6). Similarly, the highest resolution with Setting B was obtained at 60,000× magnification, which correspond to the sample scale of 0.95 Å/pixel (Fig. 6). Changing the sample scales on the detector pixel to those on the detector surface, these values are both 0.15 Å/µm (Fig. S7). It further suggests that the highest resolution is performed when the JEOL 200kV TEMs magnify the image at 0.15 Å/µm on the detector surface. The difference of the highest-resolution’s nominal magnifications of 50,000× on K2 summit and 60,000× on DE-20 is caused by the difference of the detector positions in each TEM.

**Figure 6.**
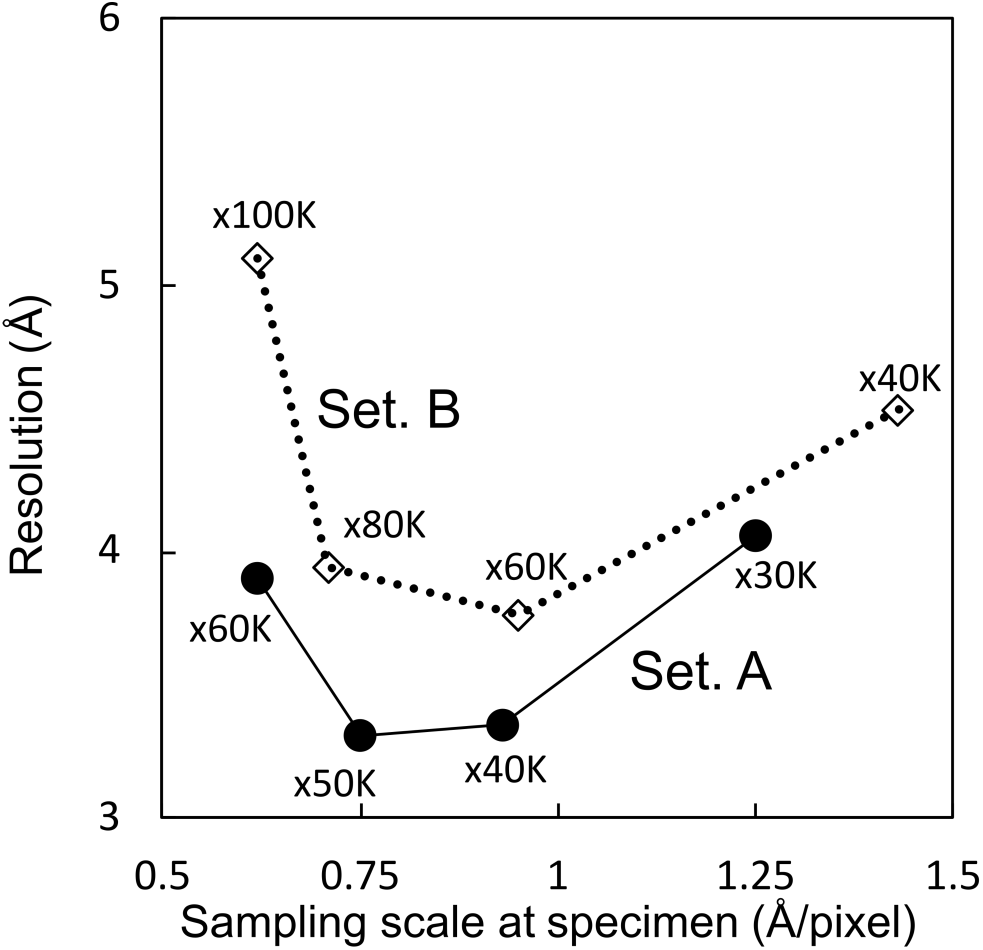
Plot of the achieved resolution of each magnification with different EM settings. The achieved resolutions of Setting A (solid dots) and Setting B (empty diamonds) at each magnification are connected by solid and dot lines, respectively. X-axis shows the sampling scale at the specimen.

When calculating Rosenthal-Henderson (“B-factor”) plots (Fig. 7), Setting A has a clear advantage over Setting B. The 50,000× Setting A dataset estimates the best B-factor (−161) with 40,000× a close second (−172). All Setting A datasets show a non-linear relationship between the points; as particle count drops, resolution decrease accelerates (Fig. 7A). Setting B demonstrates a similar non-linear relationship in datasets where the estimated B-factor is closer to those in Setting A. However, in the two datasets which estimate poor B-factor (approximately −300 or worse) a linear correlation appears (Fig. 7B).

**Figure 7.**
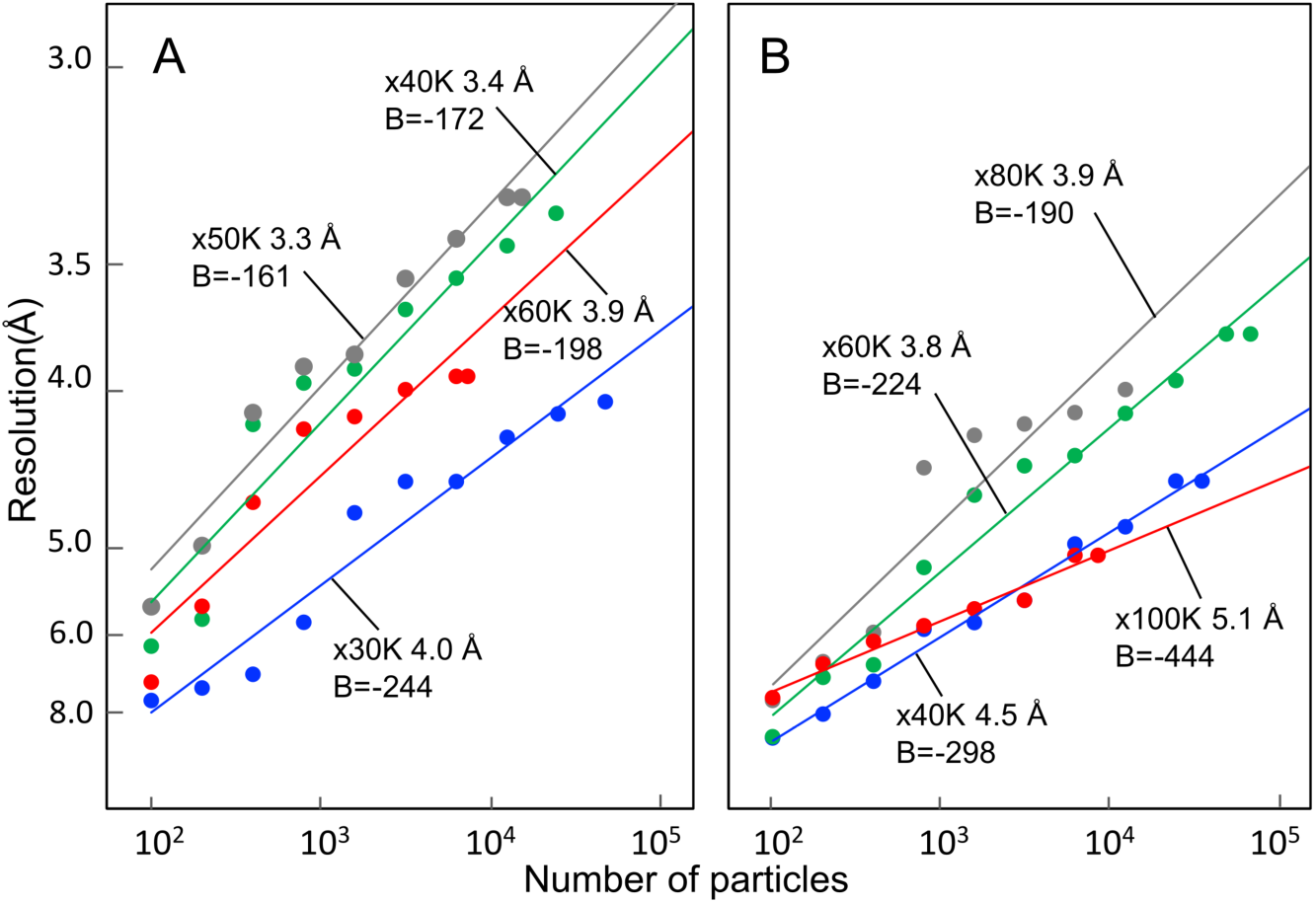
B-factor plots of all reconstructions. The points were calculated by *bfactor_plot*.*py* in RELION 3.0 at different magnifications. B-factor values are estimated from the fitted slopes. (A) Setting A and (B) Setting B. Magnifications are labelled appropriately.

### Automated acquisition

We used β-galactosidase for the sample studied with automation in multi-purpose TEM to demonstrate that usable resolutions (Fig. 8A, B) may be achieved with things other than apoferritin. Further details are listed in Table S1. In roughly the same amount of time – one half an operator day – that we were able to collect ∼150 micrographs manually, automation via SerialEM allowed acquisition of ∼450. This is without any of the more advanced automatic collection methods such as beam shift acquisition (Cheng et al., 2018). Of these micrographs, 370 were deemed usable post-motion correction and CTF estimation. With automated collection, we achieved a 3.6 Å (Fig. 8C) resolution reconstruction of β-galactosidase (Fig. 8). Calculated B-factor (−162) is comparable to those of the apoferritin limited datasets of Setting A 40,000× and 50,000× (Fig. 8D).

**Figure 8.**
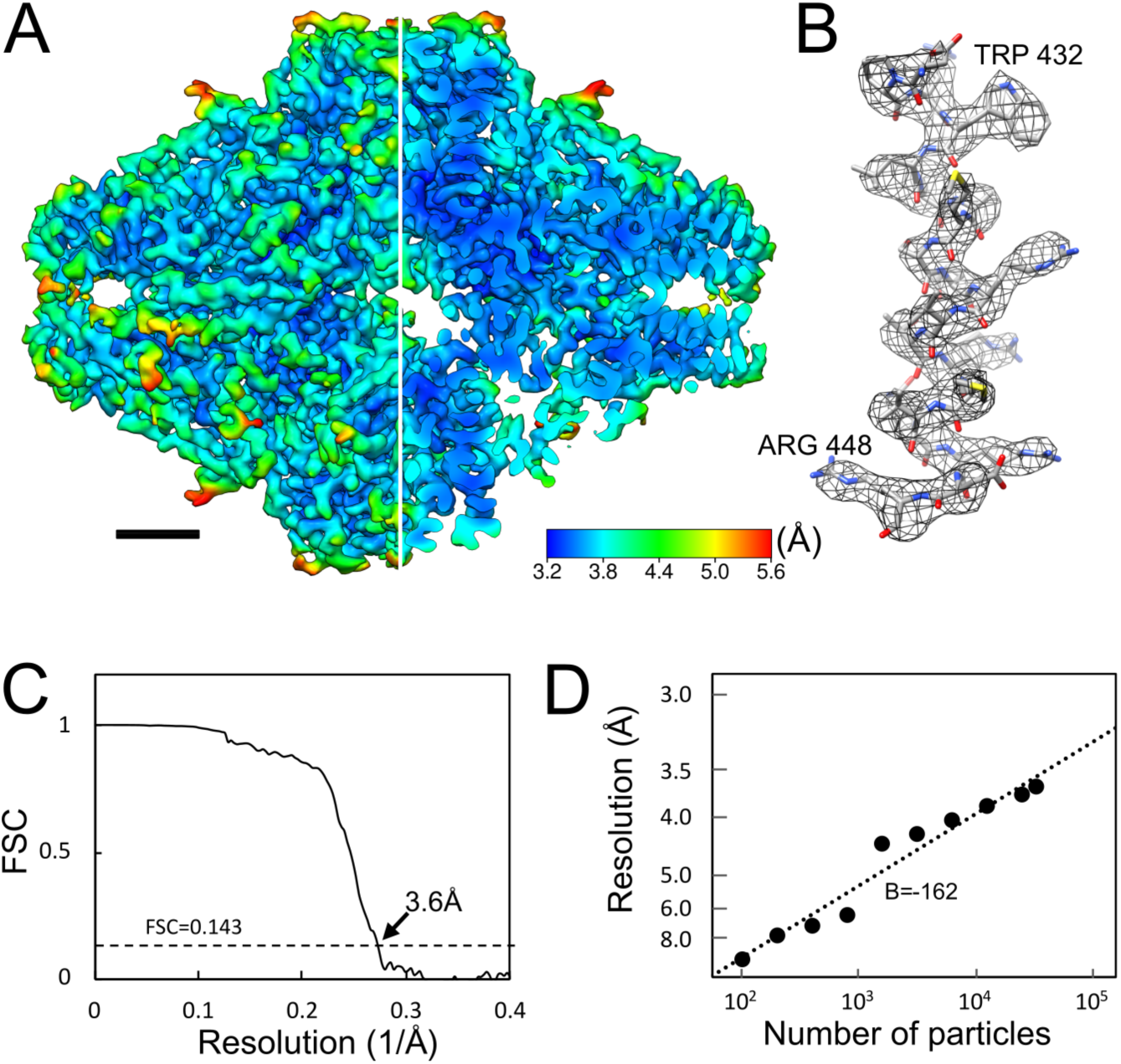
β-galactosidase reconstruction from Setting A using SerialEM automated software. Data was acquired in the same time period as other datasets, except via automation rather than manual operation. A) 3.6 Å (global estimated resolution) reconstruction coloured by local resolution, one quarter sliced away to allow visualisation of internal density. Map is contoured at 5σ. B) Representative helix (residues ASP429-ARG448) showing side-chain clarity. C) FSC curve. D) Rosenthal-Henderson plot, estimating B-factor. Estimated B-factor is dependent on whether extremely low-particle-count reconstructions are included, which influences the overall B-factor estimate.

## Discussion

After calculation of the Rosenthal-Henderson plots (Fig. 7) for all datasets, we estimated that collecting double or triple the original dataset for the Setting A at 50,000× would achieve 3 Å global resolution. We collected further 126 micrographs to complement the 153 original micrograph movies – this approximately doubled the number of usable particles but was slightly less than double the raw number of micrographs. Using RELION 3.0.8, this did not improve resolution achieved, however moving to a testing build of RELION 3.1, with improved calculation for optical parameters, permitted a 3.0 Å reconstruction with local resolution extending to 2.6 Å when calculated by *blocres* (Heymann, 2001; Heymann and Belnap, 2007). After searching the EMDB (Patwardhan, 2017) this is currently the highest resolution achieved when using a 200KV microscope and (non-autoloader) side-entry cryo-holder system.

### Comparison of EM Settings A and B

The first test of a reconstruction is the reported resolution and coincident FSC curve (Fig. S1A, Fig. 5). With this metric, and using the “limited quantity” datasets, Setting A is generally 0.5 Å superior to Setting B, except for 60,000× (Setting A) and 100,000× (Setting B), which can be attributed to the low total number of particles in the narrower field of view of the K2 Summit DED (Fig 2, Table 2). However, if comparing 60,000× (Setting A) with 100,000× (Setting B) (where both have a sampling frequency of 0.62 Å/pixel) then again Setting A becomes superior. While this absolute scale comparison is useful, 100,000× magnification for Setting B has a commensurately higher dose rate and suffers from more instability/drift than 60,000× of Setting A, which is reflected in the Rosenthal-Henderson B-factor estimation (Fig. 7B).

The best resolution was achieved at different magnifications in each setting (Figs. 4-6). When we processed the limited micrographs using RELION 3.0, the highest “limited set” resolution of 3.3 Å was obtained at 50,000× magnification with Setting A, while the highest resolution of 3.8 Å was obtained at 60,000× magnification with Setting B (Fig. 5, Table 2). Setting A would appear to have optimal conditions at 40,000-50,000×; which magnification recommended would depend on the molecular weight of the sample. Setting B showed little difference between three magnifications (60,000× and 80,000× are detailed herein, the intermediate of the three – 50,000× – is not), although the operator reported that 80,000× was more challenging to acquire than 50,000× or 60,000×, while 100,000× was even more so. Setting B provides the ability to collect superior numbers of particles from the same number of micrographs, which may prove advantageous in the case of larger or more heterogenous samples. The use of an Omega filter on Setting B does not appear to either help or hinder processing of data, although it does improve image contrast moderately (Fig. 2) which makes manual data acquisition more user friendly.

We hypothesise that the superior performance of Setting A when compared to Setting B is primarily the result of better spherical aberration (Cs: 2 mm) of the JEM-2100F microscope over the JEM-2200FS (Cs: 4.2 mm, which has a worse-than-normal Cs due to modifications necessary for the use of Zernike phase plate hardware) and improved DQE of the K2 Summit DED in the lower frequencies. It would be interesting (and troublesome) to swap the detectors between microscopes and collect further datasets to identify which of these factors has the greater effect.

### Behaviour of microscope Settings at different magnifications

When the sampling scales on each detector are adjusted respective to the detector surface area, the scale of 50,000× on K2 Summit detector (0.15 Å/µm) of Setting A corresponds to 60,000× on DE-20 detector (0.15 Å/µm) of Setting B. At this point the conditions for the best resolution were coincident between the two acquisition settings (Fig. S7). Similar achievable reconstruction resolution was demonstrated at slightly higher and lower TEM magnifications, with resolution suffering as sampling scale deviated from 0.15 Å/µm at the detector face. This would suggest that there is an optimal magnification for both combinations of microscope and detector at the same detector scale. This should be further investigated across a wider range of microscopes and detectors.

In the case of analysing with the same number of particles, it was thought that the resolution improves as the sampling scale becomes smaller. In our processing, this is untrue. This is shown by B-factor plot (Fig. 7) selecting at a given number of particles (e.g.: 10,000 total particles). At higher magnifications, the vibrations and drift that come from the side-entry cryo-holder become more critical and influence the data acquisition in a greater manner. While at lower magnifications, drift and vibrations are less of an issue, and the lower magnification becomes the limiting factor at the detector. Dose also plays a significant role in the quality of acquired data; at higher magnifications the electron dose per Å2 at the sample increases negative effects such as radiation damage, charging, and heating effects resulting in loss of sample mass coupled with large drifts as ice is vaporised. The above phenomenon occurs even though motion correction is performed. We hypothesise that adequate motion correction cannot be performed at a frame processing interval of 0.2 s (5 frames per second, fps) at higher dose rates; testing with detectors capable of outputting final micrograph movies of 20fps or greater may prove whether higher dose rates can be offset by higher framerates.

While particle polishing acts on a per-particle basis, having good drift correction across initial micrographs results in loss of fewer particles during 2D and 3D classification. In this, patch correction as implemented by MotionCor2 (or the RELION 3 implementation) is superior to whole-frame correction used by the DE-20 manufacturer scripts (Direct Electron, LP) or UNBLUR (Grant and Grigorieff, 2015), although less drift is still preferable. This improved sample stability is one of the advantages of autoloader-equipped microscopes.

Although motion correction with dose weighting was performed, there is evidence that at 60,000× (Setting A) and 100,000× (Setting B) magnifications that when the dose is optimal for a K2 Summit detector, it is too strong for the sample. Thus, there is increased motion of the sample as the vitrified ice is warmed, combined with faster sample destruction from the electron beam which together cannot be adequately compensated by per-frame motion processing. Because of the nature of recent direct electron detectors, the dose at the sample is determined by the optimal dosage for the detector; to achieve electron counting measurements, this requires a certain minimum dose per detector pixel. At higher magnifications, this can result in extremely high doses on the specimen. Stability dependence is further demonstrated by the power spectra acquired using Pt-Ir film (Fig. S4), where Setting B proves difficult to acquire high quality stable micrographs and which is reflected in the quality of the reconstruction at 100,000×. For Setting A, Thon rings are distinguishable to <2 Å for Setting A (Fig S4A, B), while the last consecutive Thon ring is visible at 2.96 Å (Fig. S4C, D) for Setting B. At lower magnifications, the limiting factor becomes Nyquist frequency and DQE. These are likely the reasons for the “sweet spot” for resolution in our datasets (Fig. 6).

### Validation of the EM-maps

While the gold-standard FSC is a simple “one number” report of a cryo-EM reconstruction, it is, more precisely, simply a measure of correlation between two independently refined half-maps (Chen et al., 2013; Herzik et al., 2019a; Scheres and Chen, 2012) and thus does not reflect local resolution variability. Map-to-model (MM) FSC is a comparison between a simulated volume generated by a fitted PDB model and either one of the two final half-maps of the GS-FSC or the post-processed (sharpened) full map. It is useful as a quality-of-fit metric for an atomic model, particularly if fitting the model via one half-map, and comparing against the second. The MM-FSC for all Setting A maps is superior to those of the Setting B maps (Fig. 5), although while apoferritin is generally rigid, there is inherent flexibility in protein which will result in minor differences between multiple datasets of the same sample.

Local resolution (recently combined with local filtering of a map) provides a finer-grain view of the quality of a reconstruction. The central core of proteins or complexes are generally higher in resolution as they show less flexibility, although this is not always true as some proteins/complexes have highly mobile active sites for substrate binding (Nakane et al., 2018). Apoferritin shows highest resolution at the interaction face between the four bundled helices of each subunit; the contact points of the subunits and the external surfaces show lower resolution. The weakest point of an apoferritin reconstruction is the N-terminal of each subunit, similar to X-ray crystallography where unstructured terminus regions are difficult to resolve.

There are many different programs for calculating local resolution; we tested several in the process of this work – although we did not carry out an exhaustive comparison of all the options now available – and finally used the Bsoft (Heymann, 2001; Heymann and Belnap, 2007) *blocres* module similar to recent work by the Lander laboratory (Herzik et al., 2017; Herzik et al., 2019a; Herzik et al., 2019b) as it estimates a range of local resolutions while neither obviously over- or under-estimating, ignoring symmetry or including dramatic resolution transitions.

Recent developments (Frenz et al., 2017; Igaev et al., 2019) in molecular dynamics programs have provided methods for fitting atomic models to electron density maps using methods independent of classical crystallographic programs such as COOT (Casanal et al., 2019; Emsley and Cowtan, 2004) or PHENIX (Adams et al., 2010), which have themselves added cryo-EM optimised functions, and which would appear to work well for “mid-to-low” resolution cryo-EM maps which have previously been difficult to interpret with the crystallographic packages. We have not used these here, although are investigating their use for other protein complexes. Ultimately, the human eye remains a good – if potentially biased – method of examining cryo-EM maps. The resolution bounds previously described (Rosenthal and Rubinstein, 2015) for the level of detail which can be expected are a good guide, and it is often immediately apparent whether a map may be the resolution it purports to be.

### Structural variability of apoferritin

The highest resolution map is that acquired at 50,000× on Setting A (Fig. 1A), where ARG63 shows two distinct conformations (Fig. 1B, marked with black arrow). This can be used as a further marker of map quality, when density for rotamers becomes distinct. While not as clear as the example of the T20S proteasome (Herzik et al., 2017), conformational shifts upon metal binding would favour increased metal coordination. Further, apoferritin does exhibit minor flexibility when reconstructed without symmetry (at a much-reduced resolution) or with symmetry expansion (Fig. S8).

Performing symmetry expansion in RELION 3.0 of 50,000× Setting A and processing with C1 symmetry (Fig. S8A) allows some flexibility in the structure (Fig S8B, resolution variations between subunits at the 3-fold symmetry site) to appear; the postprocessing-estimated B-factor value also improves slightly (from −93 to −86) although final estimated resolution of 3.3 Å was unaffected (Fig. S8C). Side chain estimated resolution decreases in an example subunit when reconstructed with symmetry expanded asymmetry (Fig. S8E) compared to when octahedral symmetry is imposed (Fig. S8D), and density for the unstructured loop (Fig. S8E, marked with black arrow) is stronger. A Rosenthal-Henderson plot was not calculated for the symmetry expanded reconstruction, although the RELION 3.1-processed increased dataset (Fig. 1) reported again a slightly higher Rosenthal-Henderson B-factor estimation (−152) than the RELION 3.0-processed 150 micrograph dataset (−161).

The behaviour of the datasets with respect to Rosenthal-Henderson plots and estimated B-factor shows a potential relationship between increased field of view and lower estimated B-factor. The lowest magnification of Setting A (30,000×) estimates similar B-factors to Setting B (60,000× and 80,000×), which has a much wider field of view (Fig. 2) although the very high magnifications show similarly poor B-factor estimates (Fig. 7). We attribute these effects to microscope, cryo-holder stability, and radiation damage at the higher magnifications and to increased area for local motions to occur at lower magnifications.

### Applicability of multi-purpose microscopes in cryo-EM SPA

How valid is the use of a multi-purpose TEM for SPA? It will not challenge the most expensive equipment combinations; however, this work demonstrates that even when imposing a limit on data quantity, it is possible to achieve usable resolutions for *de novo* structure determination, provided that other biochemical data is known. For the purpose, it is important to understand the limits created by equipment: which magnifications provide the least drift, most easily optimised dose rate; the detector will influence the rate at which data can be acquired, and potentially usable particles per micrograph. At these resolutions, DQE appears to have less of an impact than was hypothesised.

When adding automated acquisition, quantity of data acquired during a similar time period increases, but percentage of good micrographs drops slightly when compared to a skilled operator. Although the potential advantages for users with respect to use of time and work environment cannot be overstated, it remains necessary to maintain regular vigilance of cryogen levels as they are not automatically replenished. The lower symmetry of β-galactosidase offsets the increased quantity of data that was collected using SerialEM automation software (Fig. 8), although in terms of particles picked the two proteins were broadly similar (Table 2, Table S1) and demonstrates that acquisition of lower symmetry protein complexes is viable to achieve usable datasets.

There is some utility in comparison of our results to far superior ones of the same complex obtained elsewhere, such as EMD-9599 (Danev et al., 2019), EMD-9865 (Kato et al., 2019) in the electron microscopy data bank (EMDB), as they were acquired using equipment providing the very highest performance possible for cryo-EM SPA. The 1.75 Å map of apoferritin (Wu et al., 2020) from a 200KV microscope further demonstrates the possibilities with increased quantities of data when coupled to optimal grids, sample preparation technique and microscope stability. Although of far greater value is comparing against other datasets, such as the T20S proteasome (Herzik et al., 2017) where ∼3 Å (2.8 Å highest local resolution) allowed identification of side chain orientation and tightly bound water molecules on a 200kV microscope, albeit one equipped with an autoloader stage and optimised for SPA. We achieved similar clarity with respect to potential metal-binding sites and residues showing different rotamer states (Fig. 1B, C). While the symmetry of T20S proteasome is not as high (D7, or 14-fold) as apoferritin, the number of particles required for this was an order of magnitude higher than the quantity of data acquired herein, however, exceeding 1,000,000 particles picked and ∼80,000 used in the final reconstruction.

The recent work demonstrating the application of conventional 200kV hardware to proteins <100kDa (Herzik et al., 2019b) indicates that phase-contrast systems (Danev et al., 2014) are not a requirement for analysis of proteins of that approximate mass. Herzik *et al*. do further discuss the difficulties in processing a 50kDa complex, concluding that phase-contrast may be necessary for these very small complexes, although clarity in 2D class averages may indicate that improvements in image processing solutions would be sufficient. With the improvements demonstrated here when reprocessing a dataset in RELION 3.1 versus RELION 3.0, we agree.

For SPA using a multi-purpose TEM, the two primary limiting factors are the ability to collect large volumes of data and maintain sample quality – the latter being something which also affects every TEM, and therefore we consider a null concern for this situation. By achieving a global resolution of 3 Å with a multi-purpose TEM with widely available hardware and limited datasets and comparing two multi-purpose TEM setups, we hope that it will encourage potential users of cryo-EM SPA who do not have access to state-of-the-art facilities.

## Acknowledgements

The authors thank Dr. H. Yanagisawa for the gift of the apoferritin test sample, Dr. Jaap Brink (JEOL USA) and Dr. David Mastronarde for help setting up the SerialEM software, and Mr. Yoshihiro Arai (Terabase Inc.) and Mr. Takayuki Owada (Kyodo Printing Co. Ltd.) for renting a Gatan K2 Summit DED. The study was supported by a collaboration program with Terabase Inc.\

## Data availability

The cryo-EM reconstructions of all post-processed maps and half maps have been deposited in the EMDB under the accession codes: EMD-30096 (3 Å Setting A 50,000×, RELION 3.1), EMD-30101 (Setting A, 30,000×, RELION 3.0), EMD-30100 (Setting A, 40,000×, RELION 3.0), EMD-30098 (Setting A, 50,000×, RELION 3.0), EMD-30099 (Setting A, 60,000×, RELION 3.0), EMD-30097 (Setting A, 50,000×, RELION 3.0, symmetry expanded), EMD-30103 (Setting B, 40,000×, RELION 3.0), EMD-30105 (Setting B, 60,000×, RELION 3.0), EMD-30106 (Setting B, 80,000×, RELION 3.0), EMD-30107 (Setting B, 100,000×, RELION 3.0), EMD-30095 (Setting A, 40,000×, RELION 3.0, Serial-EM acquisition, β-galactosidase).

## Author contributions

Sample preparation: TK, CS, NT, data acquisition; YK, CS, RNBS, KM; data processing; RNBS, YK, KM, automatic data acquisition: TK, CS, RNBS, KM, draft manuscript; RNBS, YK, manuscript revision; RNBS, YK, CS, KM, conceptualisation; KM, project oversight; KM.

## Methods

### Cryo-electron microscopy

High-symmetry (24-fold/octahedral) heavy-chain apoferritin was used as a test specimen, which was the generous gift of Dr. H. Yanagisawa, University of Tokyo. β-galactosidase (SIGMA-ALDRICH, St. Louis, MO) was purified by gel filtration on a Superdex-200 size-exclusion chromatography column connected to an ÄKTA FPLC apparatus (GE Healthcare Bio-Sciences, Piscataway, NJ) with an elution buffer comprised of 25 mM Tris (pH 8), 50 mM NaCl, 2 Mm MgCl_2_ and 1mM TCEP. An aliquot of sample solution was applied to standard molybdenum Quantifoil grids R1.2/1.3 (Quantifoil Micro Tools GmbH) and vitrified by rapid plunging in liquefied ethane using a Vitrobot Mark IV (Thermo Fisher Scientific) at 95% humidity and 4 °C. The frozen grid was mounted on a Gatan 626 cryo-transfer specimen holder at liquid nitrogen temperature and loaded into either a JEM2100F microscope (JEOL) equipped with a K2 Summit DED (Gatan) (Setting A) or a JEM2200FS microscope (JEOL) equipped with a DE-20 DED (Direct Electron LP) (Setting B). Magnifications were varied and are detailed in Table 2. Both microscopes were operated with thermal Schottky electron source at 200kV. In the case of JEM2200FS, an Omega-type energy filter was used with a slit width of 15 eV. Spherical aberrations of each pole piece were 2.0 mm (JEM2100F) and 4.2 mm (JEM2200FS). The illumination conditions were optimized for K2 Summit counting mode (8 e-/pixel/sec on detector) and maintained through DE-20 acquisition for comparison purposes via low dose acquisition, although at 80,000× and 100,000× magnification the dose rate for DE-20 acquired data was considered very high. Further details can be found in Table 2. Movies were collected over a minimum of 3 seconds at 5 fps (frames per second) in both detectors.

### Image processing

Movies were motion-corrected using the MotionCor2 algorithm (Zheng et al., 2017) as implemented in RELION 3 (Zivanov et al., 2018) using dose-weighting and patch correction with 5×5 grid for K2 Summit DED and 5×3 grid for the DE-20 DED. CTF was estimated by CTFFIND4 (4.1.10) (Rohou and Grigorieff, 2015). Particles were picked by either a) using the RELION 3 LoG-based (Laplacian of Gaussian) auto picker (Zivanov et al., 2018) or b) manually picking 100-200 particles, as the shape of apoferritin means that more particles are not needed for autopicking. The box size was determined so that one edge was between 180 to 240 Å, and extracted particles were 2D-classified. Good classes were used as a reference for a second round of auto picking. The extracted particles were sorted against the autopicking references and the worst particles discarded, after which they were subjected to 3D classification using a map generated *ab initio* using the cisTEM (Grant et al., 2018) algorithm as a reference, with octahedral symmetry (Fig. S5). The best class(es) were selected, and a map was refined via Refine3D followed by postprocessing. For further optimization, CTF refinement and Bayesian polishing were performed on the obtained input particles, and Refine3D was performed again to obtain the final map. Local resolution was calculated using *blocres* (with default settings but defining symmetry) from the Bsoft package (Heymann and Belnap, 2007). The procedure is summarized in Fig. 3. Processing in RELION 3.1 proceeded in a similar fashion except for the particle sorting step, which has been removed in the current testing versions of RELION 3.1. Visualisation of 2D and 3D images were carried out using RELION (Zivanov et al., 2018), Fiji (Schindelin et al., 2012) or UCSF Chimera 1.11.2 (Pettersen et al., 2004) depending on dimensionality.

### Model fitting and map validation

An x-ray crystallographic-derived atomic model (PDBID: 2CIH) was fitted to the obtained eight maps using the “fit in map” function in UCSF Chimera (1.11.2) (Pettersen et al. 2004). The maps were segmented using SEGGER (v1.4.9) (Pintilie et al., 2010) and an α-helix corresponding to residues 48-77 was visualized for each map. FSC curves of each of the data sets were calculated. The correlation between half-maps (GS-FSC) (Chen et al., 2013) was estimated by 0.143 cut-off (Rosenthal and Henderson, 2003), and the correlation between map and atomic model (PDB ID: 2CIH) (MM-FSC) was calculated with 0.5 cut-off. The PDB-based atomic model map was generated using UCSF Chimera’s “*molmap*” function (Pettersen et al., 2004) at the GS-FSC resolution of each reconstructed map. The CTFFIND estimated resolution was assessed by analysis of the logfiles generated by CTFFIND4 (Rohou and Grigorieff, 2015). B-factor plots were calculated by running an appropriately modified copy of RELION’s script, *bfactor_plot*.*py*. The script randomly selects subsets of particles from each data set and executes Refine3D and Postprocess steps. Plotting the natural logarithm of each particle subset against the inverse of the squared resolution for each refinement allows estimation of particle and dataset quality by correlation of a linear fit (Zivanov et al., 2018).

### Assessment of TEM’s performance

DQE curves were measured using the shadow of a beam stopper. The obtained data was processed by FindDQE (Ruskin et al., 2013). Thon rings are compared with two TEM settings by using FFT of Pt-Ir micrograms with the same data acquisition condition. Radial profiles were generated by Gatan Microscopy Suite 3 (Gatan, Inc.).

## Supplementary information

**Table S1.**
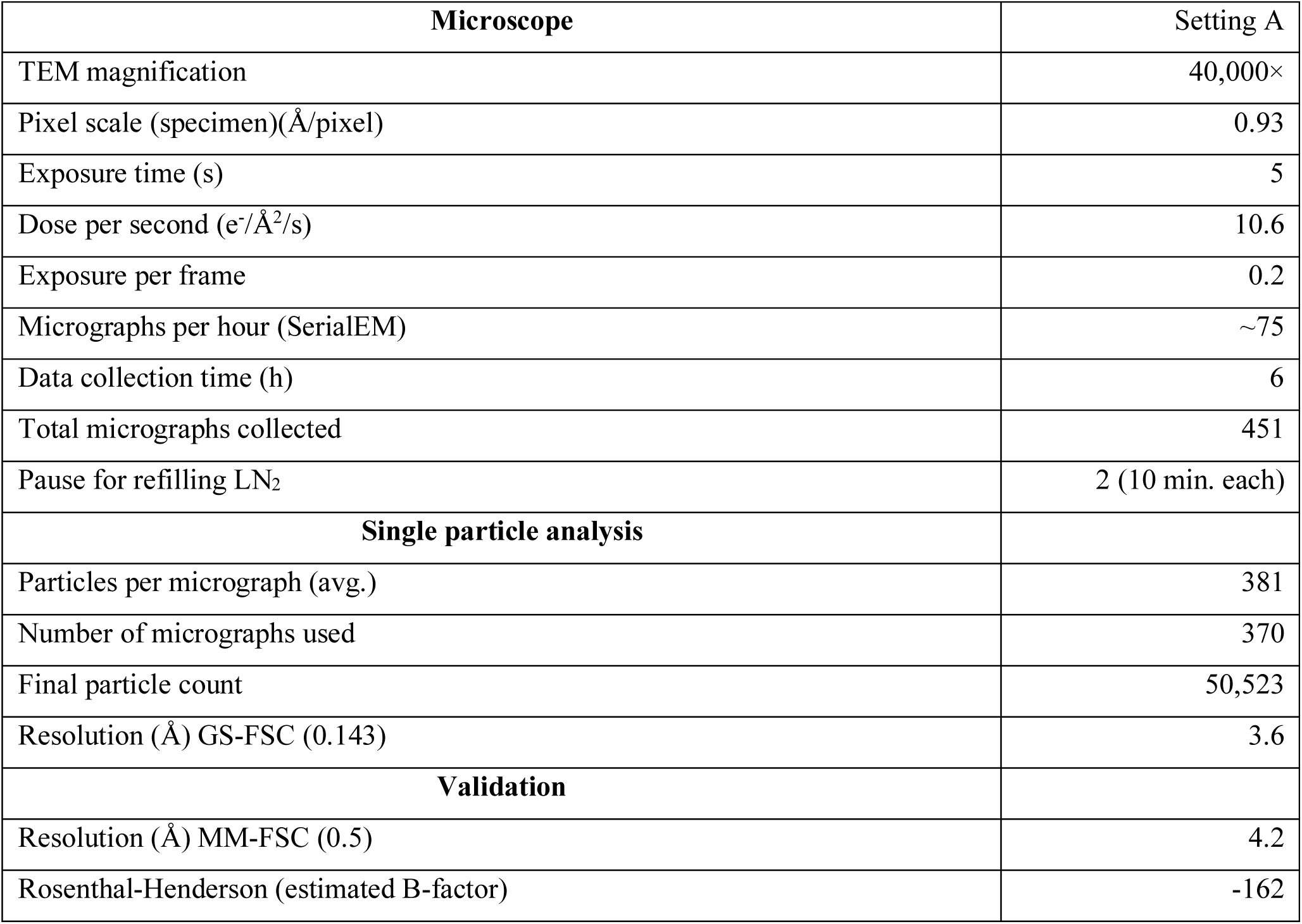
Details of acquisition conditions for β-galactosidase via SerialEM.

**Figure S1.**
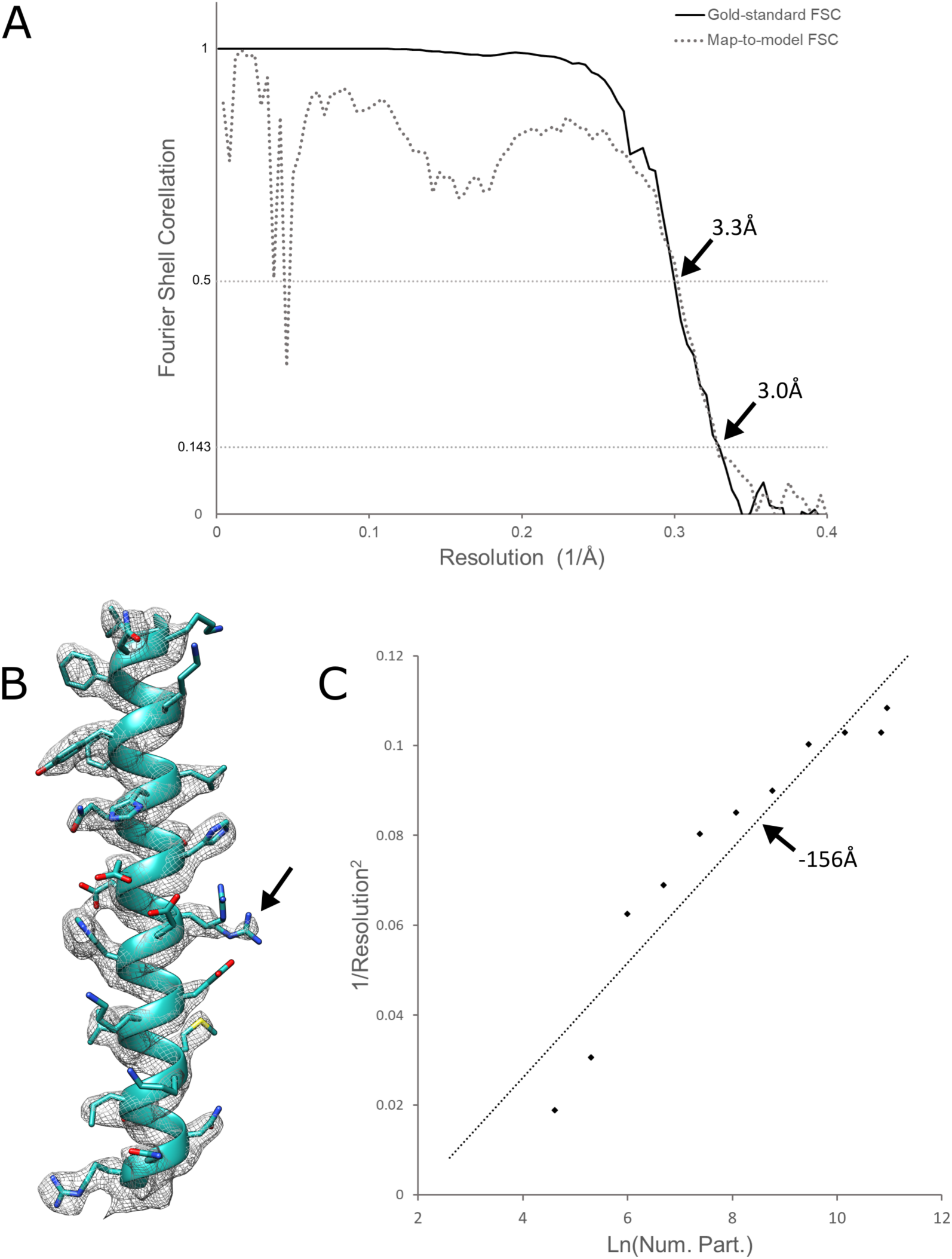
Additional data for Fig. 1. A) Gold-standard FSC (black line) and Map-to-model FSC (dashed line) for 3 Å (global estimate) apoferritin reconstruction. B) The helix from Fig. 1B, contoured at 5σ rather than 3σ (Fig. 1B), permitting visualisation of the loss of one of the ARG63 rotamer densities, indicating that it may be a less favourable conformation. C) Rosenthal-Henderson plot, estimating B-factor to be −162.

**Figure S2.**
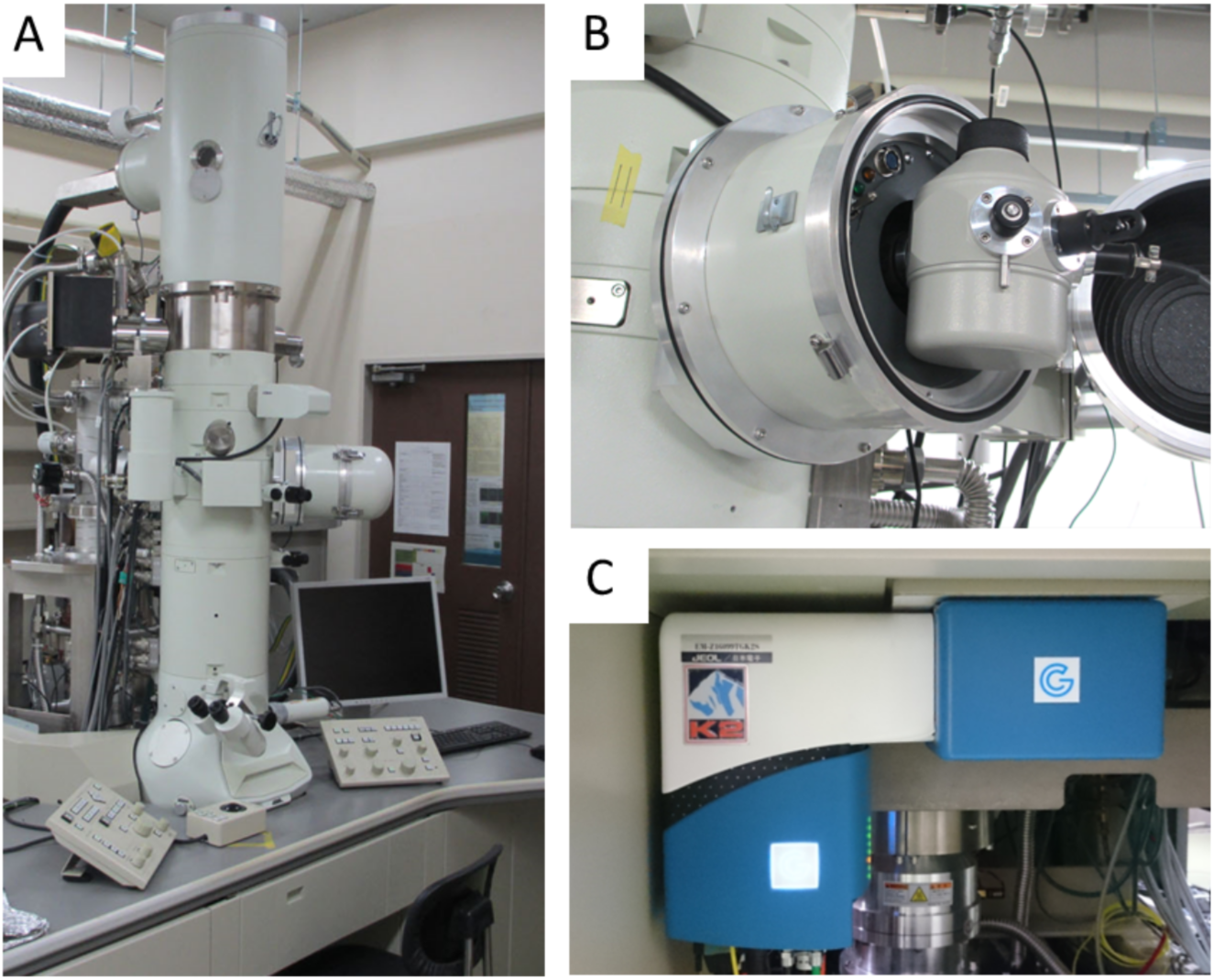
One of two electron microscope settings. Gatan K2 Summit DED (C) is mounted on JEM-2100F microscope (A). Gatan 626 cryo-specimen holder is used to sustain the frozen grid at liquid Nitrogen temperature (B). The second electron microscope setting has been detailed previously (Murata and Wolf, 2018).

**Figure S3.**
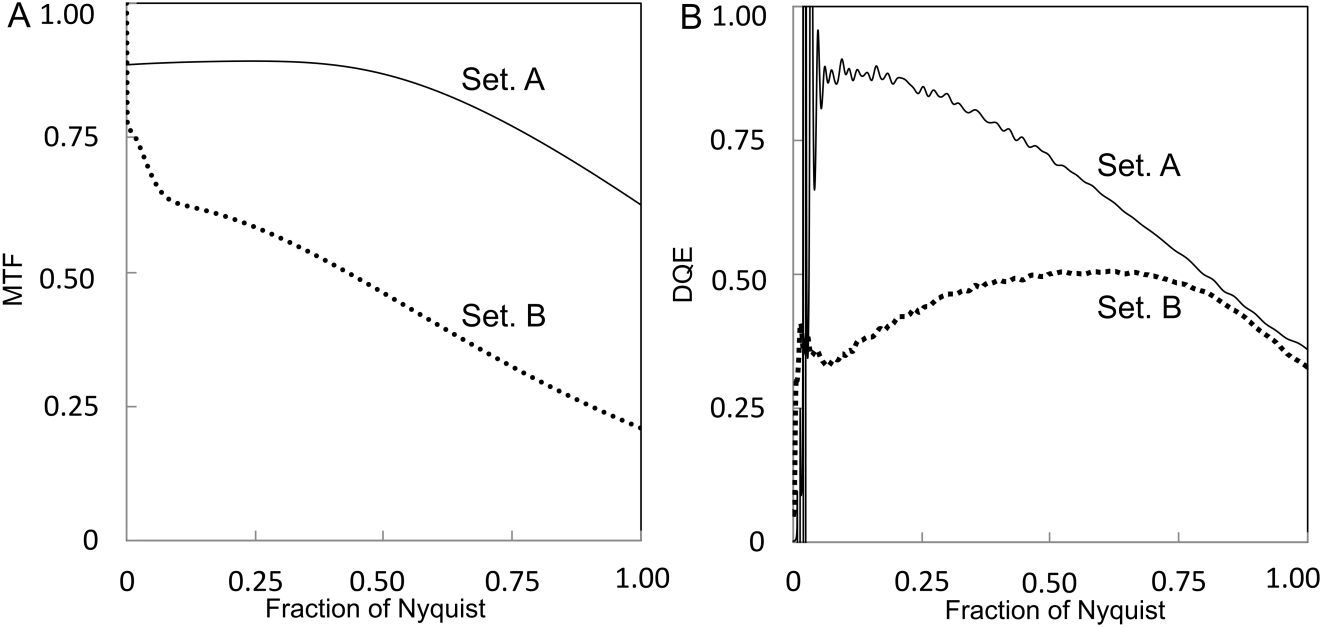
Modulation transfer function (MTF) and Detective quantum efficiency (DQE) curves in EM settings A and B. A) MTF curves for each setting, B) calculated DQE for each setting. DQE curves are estimated with a beam stopper using FindDQE software (Ruskin et al., 2013).

**Figure S4.**
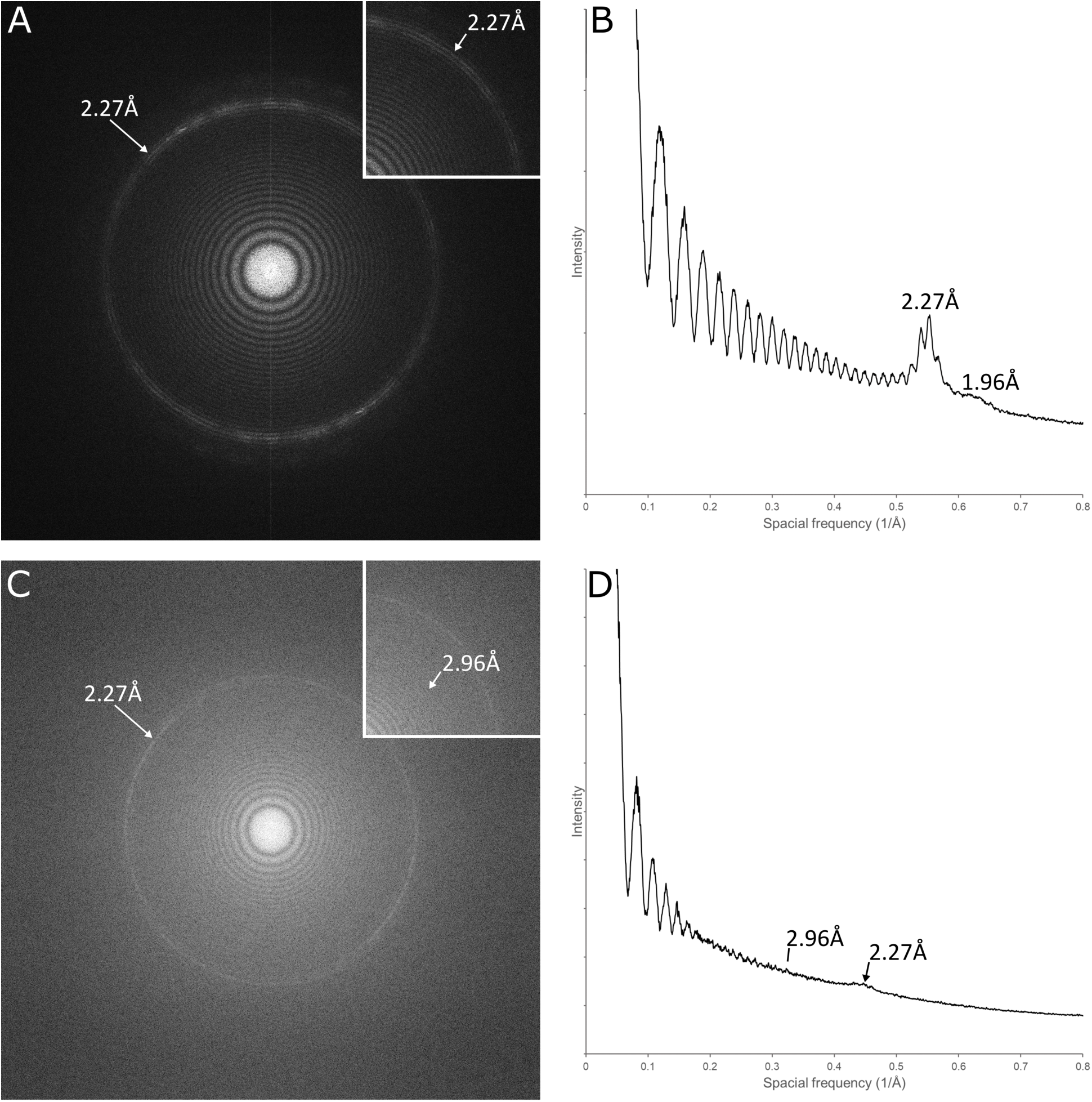
Thons rings of FFT images of Pt-Ir film. Micrographs of Pt-Ir film were acquired by Setting A and B at 100,000×, 0.5µm defocus, and the power spectrum were generated by FFT. A) Setting A power spectrum, B) Plot of rotationally averaged radial profile of (A), C) Setting B power spectrum, D) Plot of rotationally averaged radial profile of (C). With Setting A, Thon rings are clearly distinguishable to the diffraction ring; with Setting B, difficulties in maintaining stability have caused a slight drift in defocus and minor astigmatism resulting in blurring of the rotationally averaged profile.

**Figure S5.**
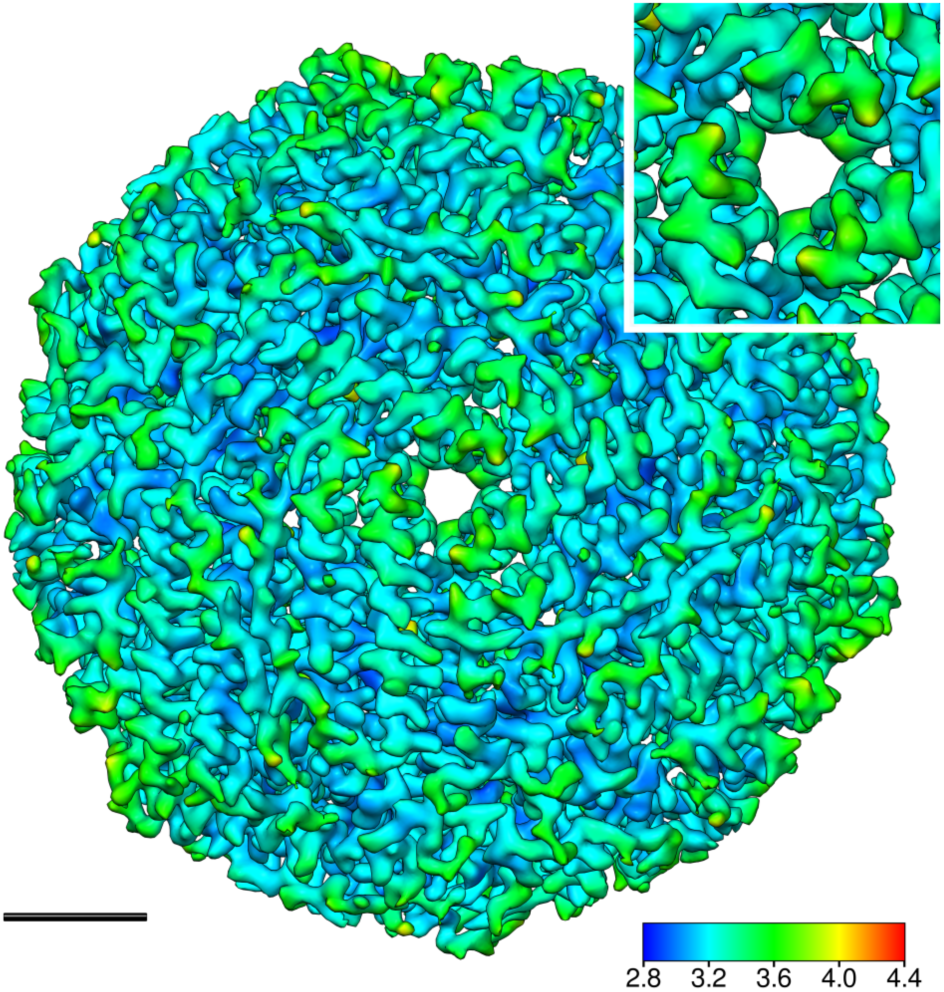
The best resolution map of apoferritin at 3.3 Å generated by Setting A. It was achieved at 50,000× magnification. The local resolution was coloured from 2.8 to 3.6 Å resolution. Surface depicted at 4σ. Breakout focussed on 3-fold symmetry axis, where estimated local resolution for each subunit is identical. Scale bar equals 2 nm.

**Figure S6.**
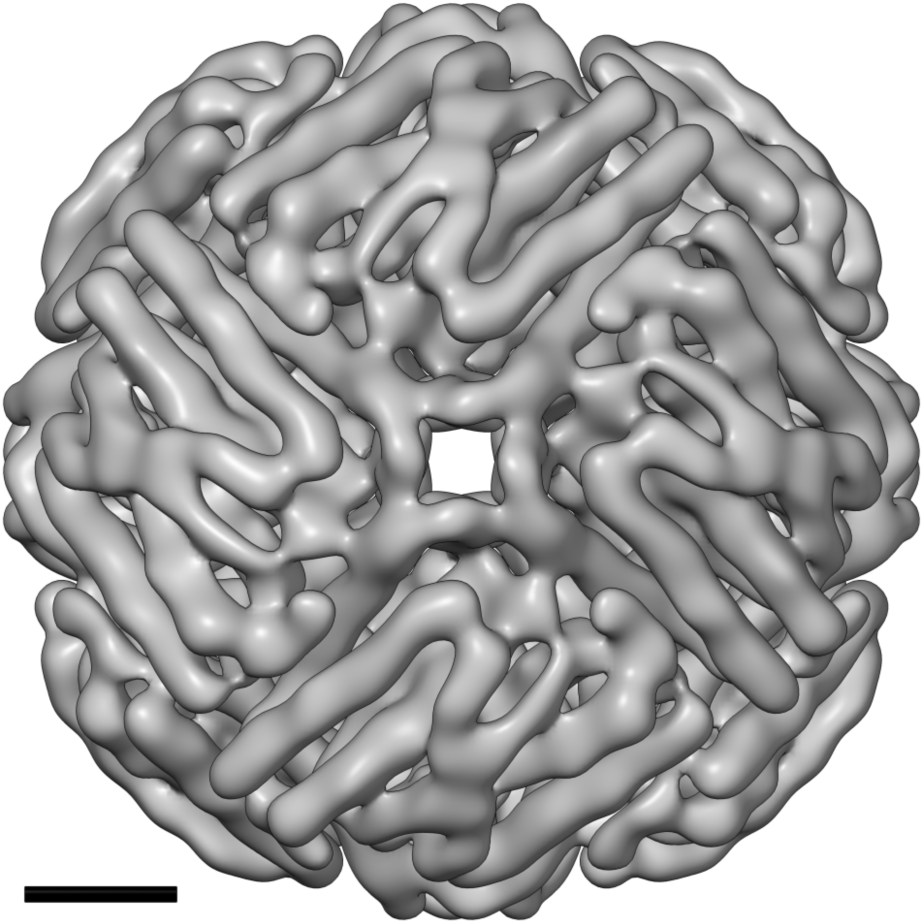
A representative *ab initio* 8 Å initial model of apoferritin. The map was generated with *cis*TEM (Grant et al., 2018). Scale bar equals 2 nm.

**Figure S7.**
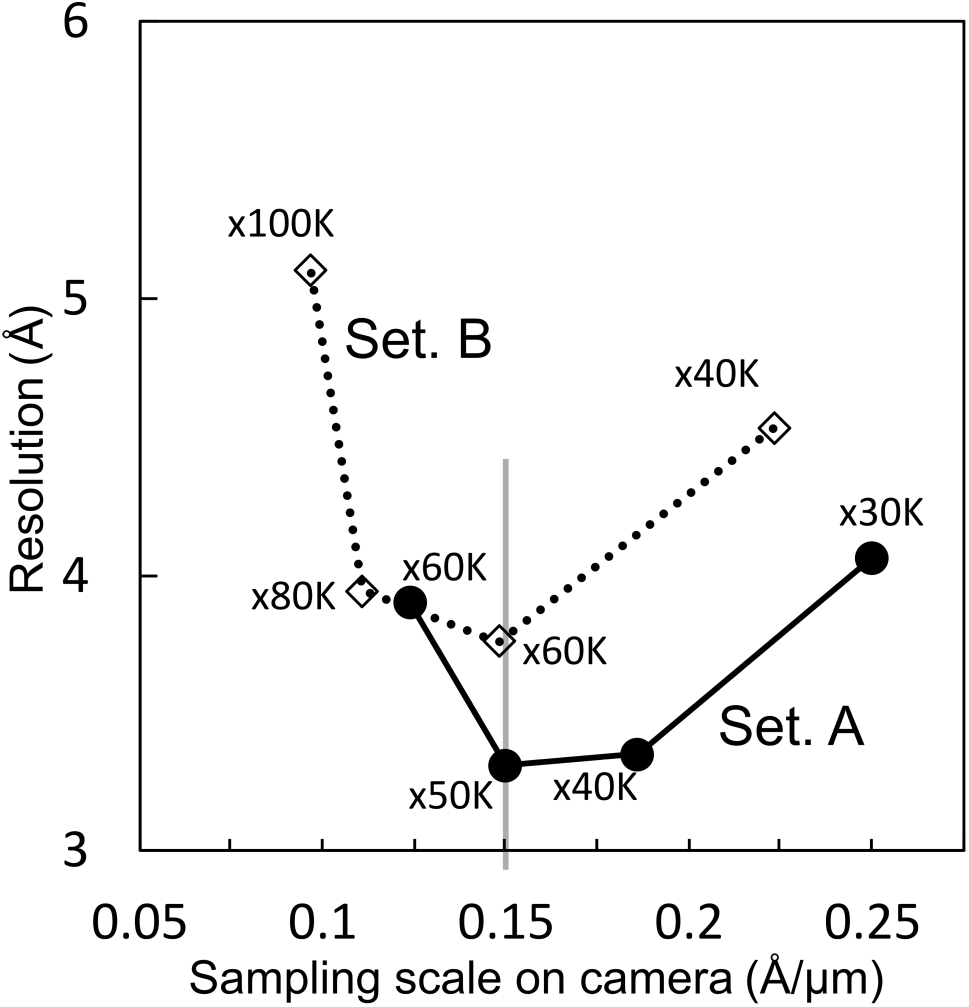
Resolution plots for sampling scales at detector face. Final global estimated resolution of each reconstruction at different magnifications was generated using Settings A and B. Vertical grey line indicates the scaling point at which maximum resolution was achieved for both Settings A and B.

**Figure S8.**
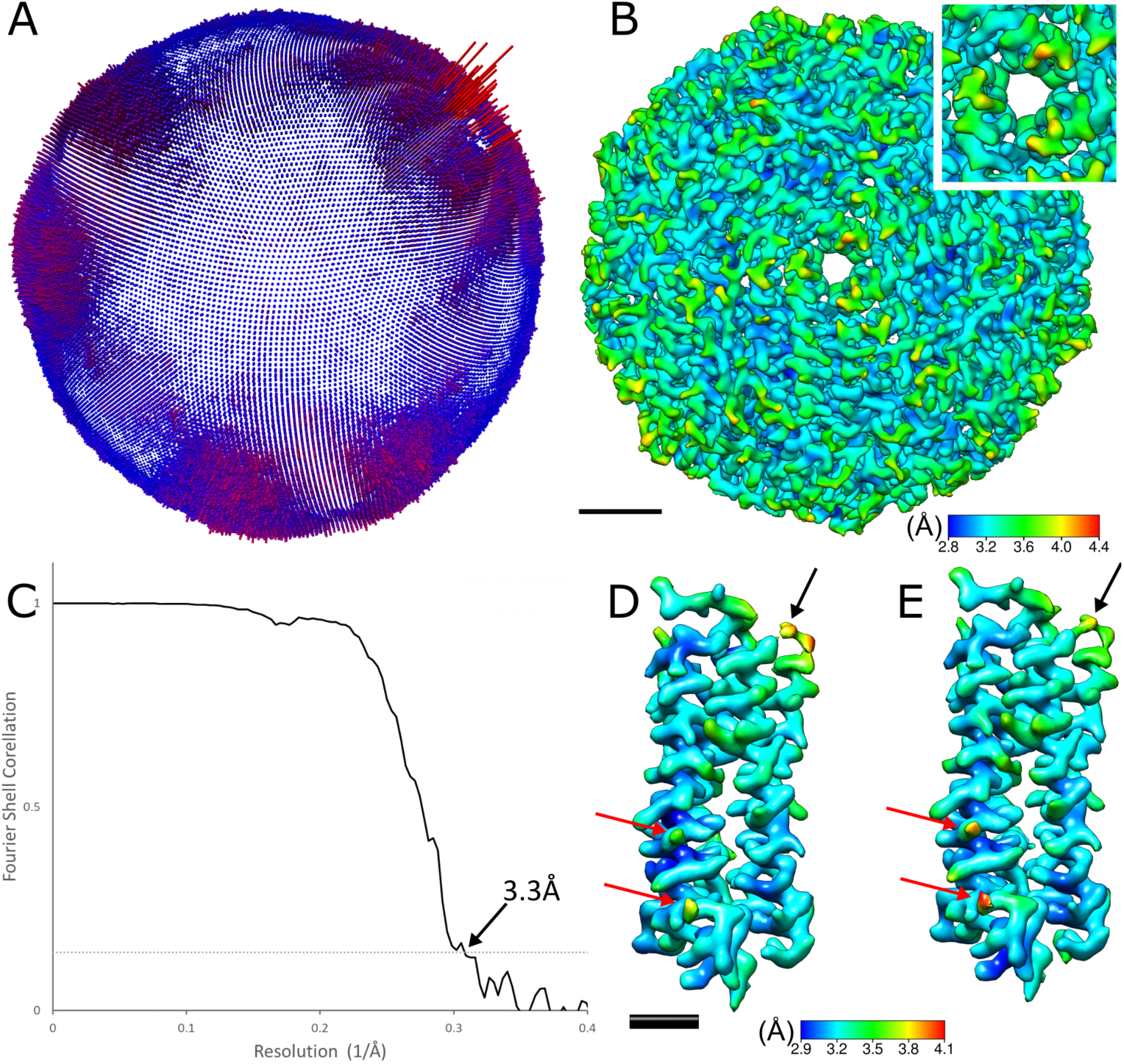
Symmetry expanded reconstruction based upon Setting A 50,000× limited dataset. The map shows variance in angles assigned, which indicates some asymmetry or flexibility in the protein complex. A) Symmetry expanded angle assignments of apoferritin reconstruction, showing deviation in angular assignment from octahedral symmetry, B) apoferritin reconstruction viewed from same angle as (A) coloured by local resolution observed from the 3-fold symmetry axis, which shows allows example visualisation of resolution variance between three subunits (focussed view in breakout). Scale bars equal 2 nm. C) FSC curve of symmetry expanded apoferritin reconstruction showing 3.3 Å resolution at GS-FSC (0.143). D) extracted subunit from symmetric reconstruction, coloured by local resolution, E) extracted subunit from symmetry expanded (asymmetric) reconstruction, coloured by local resolution. The flexible trans-helix backbone (black arrows in D, E) is contiguous and higher resolution in the asymmetric reconstruction. Some residue sidechains are lower resolution, however (red arrows in D, E). Scale bar equals 1nm.

